# MDGAs are fast-diffusing molecules that delay excitatory synapse development by altering neuroligin behavior

**DOI:** 10.1101/2021.03.16.435652

**Authors:** Andrea Toledo, Giorgia Bimbi, Mathieu Letellier, Béatrice Tessier, Sophie Daburon, Alexandre Favereaux, Ingrid Chamma, Kristel M. Vennekens, Jeroen Vanderlinden, Matthieu Sainlos, Joris de Wit, Daniel Choquet, Olivier Thoumine

## Abstract

MDGAs are molecules that can bind neuroligins *in cis* and interfere with trans-synaptic neurexin-neuroligin interactions, thereby impairing synapse development. However, the sub-cellular localization and dynamics of MDGAs, as well as their specific mode of action in neurons are still unclear. Here, using both surface immunostaining of endogenous MDGAs and single molecule tracking of recombinant MDGAs in dissociated hippocampal neurons, we show that MDGA1 and MDGA2 molecules are homogeneously distributed and exhibit fast membrane diffusion, with a small reduction in mobility across neuronal maturation in culture Using shRNAs and CRISPR/Cas9 strategies to knock-down/out MDGA1 or MDGA2, we demonstrate an increase in the density of excitatory synapses accompanied by enhanced membrane immobilization and an increase in the phosphotyrosine level of neuroligins associated with excitatory post-synaptic differentiation. Finally, we show that decreasing MDGA expression level reduces the mobility of AMPA receptors and increases the frequency of AMPA receptor mediated mEPSCs. Overall, our results support a mechanism by which interactions between MDGAs and neuroligin-1 delays the assembly of functional excitatory synapses containing AMPA receptors.

## Introduction

During brain development, synapse assembly and maturation are critical processes involving several families of adhesion molecules, among which the neurexin-neuroligin complex has been one of the most extensively studied ^1–5^. Neuroligins (NLGNs) are post-synaptic proteins that comprise four members in rodents (NLGN1-4), NLGN1 being primarily localized at excitatory synapses, NLGN2 and NLGN4 at inhibitory synapses, and NLGN3 at both types of synapses ^6, 7^. At the structural level, NLGNs form both homo- and hetero-dimers through *cis*-interactions between their acetylcholinesterase (AchE)-like domains ^8–12^. NLGNs also contain a stalk region that can be cleaved by proteases ^13, 14^, a single pass transmembrane domain, and a conserved intracellular tail whose binding to post-synaptic scaffolding molecules can be modulated by phosphorylation and thereby influence AMPA receptor recruitment ^3, 15–20^. Post-synaptic NLGNs bind pre-synaptic neurexins (NRXNs) through extracellular interactions, thus making a bridge between sub-micron adhesive modules across the synaptic cleft that precisely position AMPA receptors ^21–23^.

Apart from NRXNs, few direct NLGN binding partners have been identified ^2^. Recently, MAM-domain GPI-anchored molecules (MDGAs) were reported to bind *in cis* to NLGNs with high affinity and compete with their binding to NRXNs ^24^. In the co-culture assay, the expression of MDGAs together with NLGNs in heterologous cells impairs the synapse-inducing activity of NLGNs on contacting axons ^25–27^. Crystal structures of MDGA-NLGN complexes revealed that MDGAs bind through their first Ig1-Ig2 domains to the two lobes of the NLGN extracellular dimer, at sites which overlap with the NRXN binding interface ^25, 28, 29^. Manipulations of MDGA1 protein levels by over-expression (OE), knock-down (KD), or knock-out (KO) in neurons have led to the common view that MDGA1 selectively inhibits inhibitory synapse formation by primarily repressing NLGN2-NRXN interactions ^26, 27, 30, 31^. Similar studies performed on MDGA2 have led to more debated results, i.e. some reports showing that MDGA2 KO specifically inhibits excitatory synapse formation *in vivo* ^32^, while others showing no effect of MDGA2 KD on either excitatory or inhibitory synapses in culture ^31^.

Notwithstanding the foregoing, there are still a number of critical questions that need to be answered in order to get a more complete picture of the role of MDGAs in synapse differentiation and function ^24, 33^. 1/ What is the surface dynamics and nanoscale localization of endogenous MDGAs at the neuronal membrane? Indeed, in the absence of highly specific and efficient antibodies to MDGAs allowing reliable immunostaining in tissue, over-expression approaches have yielded contrasting results about the presence of MDGA1 and MDGA2 at excitatory versus inhibitory synapses ^26, 31^. Given the absence of an intracellular domain with potential synaptic retention motifs, MDGAs are in fact expected to display fast membrane diffusion and not accumulate at synapses. 2/ Considering that the binding of MDGAs and NRXNs to NLGNs is mutually exclusive, what is the effect of MDGAs on the dynamic distribution of NLGN in dendrites and on the NLGN-dependent phosphotyrosine signaling pathway known to regulate post-synaptic differentiation ^3, 16^? 3/ Consequently, what is the impact of MDGAs on AMPA receptor surface dynamics and synaptic function, which were shown to be tightly regulated by NLGN1 ^18, 22, 34, 35^?

To address those questions, we examined the surface localization and dynamics of MDGAs, using both custom-made antibodies to endogenous MDGA1 as well as replacement strategies with recombinant MDGAs bearing small tags and labelled with monomeric fluorescent probes. Using a combination of single molecule imaging and electrophysiology, we also assessed the effects of single-cell MDGA knock-down or knock-out on NLGN1 and AMPA receptor membrane diffusion and localization, in relation to synapse maturation. Finally, we examined the impact of MDGA knock-down on endogenous NLGN phosphotyrosine levels by biochemistry. Together, our data indicate that MDGAs are highly mobile and homogeneously distributed molecules, that alter both NLGN1 and AMPA receptor dynamics, localization, and function, thereby significantly delaying differentiation of the post-synapse.

## Results

### Endogenous MDGA1 is homogeneously distributed in hippocampal pyramidal cells

To examine the localization of endogenous MDGAs in neurons, we produced and purified full length recombinant Fc-tagged MDGA1 and MDGA2, and custom-ordered the generation of rabbit polyclonal antibodies against those proteins. We then characterized the collected antisera using immunohistochemistry and Western blots. The MDGA1 antiserum recognized recombinant HA-MDGA1 (but not HA-MDGA2) extracted from HEK293T cells at the expected molecular weight of 130 kDa on immunoblots (Figure 1A), and reactivity to MDGA1 was abolished by pre-incubation of the antiserum with an excess of recombinant MDGA1-Fc antigen (Figure 1B). The MDGA1 antiserum recognized a single band around 130 kDa in brain homogenates from wild type mice, which was not present in brain homogenates from *Mdga1* KO mice, demonstrating antibody specificity (Figure 1C). Immunohistochemistry on brain sections showed abundant MDGA1 localization in the hippocampus, with prominent labeling in CA3 and CA1 stratum radiatum and stratum oriens containing pyramidal neuron dendrites (Figure 1D). MDGA1 staining was absent in brain sections from *Mdga1* KO mice ^36^, showing antibody specificity in tissue. MDGA1 was detected both in pre-synaptic and post-synaptic fractions from synaptosome preparations, revealing its presence in synaptic compartments (Figure 1E). Unfortunately, the MDGA2 antiserum was not specific enough to be used further. However, we detected abundant levels of MDGA2 mRNAs by RT-qPCR in hippocampal cultures (Supplementary Fig. 1B), in agreement with previous *in situ* hybridization and β-galactosidase staining ^27, 32^, together suggesting that the MDGA2 protein is also expressed.

**Figure 1.**
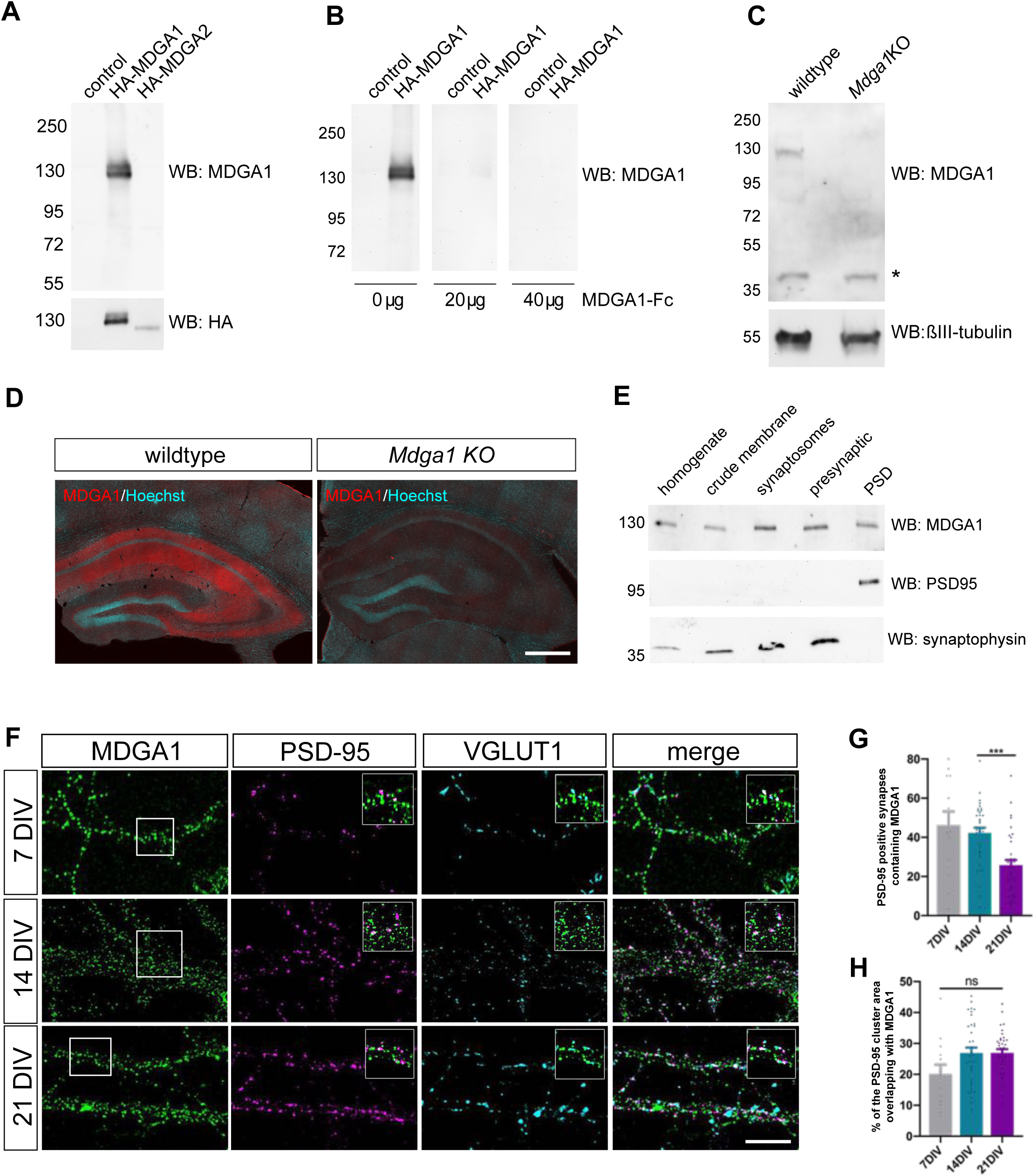
Validation of MDGA1 antibody and distribution of endogenous MDGA1 in brain slices and dissociated hippocampal cultures. **(A)** MDGA1 antiserum recognizes recombinant HA-MDGA1, but not HA-MDGA2, transiently expressed in HEK293T cells (top membrane). Mock-transfected HEK 293T cells were used as control. HA antibody labels both HA-MDGA1 and HA-MDGA2 (bottom membrane). Molecular weight markers in kDa indicated on the left. **(B)** Competition with different amounts (0, 20 and 40 µg) of excess recombinant MDGA1-Fc blocks detection of HA-MDGA1 by MDGA1 antiserum. **(C)** MDGA1 antiserum detects a single 130 kDa band in brain homogenate from wild type mice, which was absent in brain homogenate from *Mdga1* KO mice (top membrane). Asterisk indicates non-specific band. ßIII-tubulin was used as loading control (bottom membrane). **(D)** Immunohistochemistry with MDGA1 antiserum (red) reveals strong immunoreactivity in CA3 and CA1 regions of the hippocampus in wild type adult mice, which was absent in *Mdga1* KO mice. Nuclear marker Hoechst (cyan) was used to visualize tissue architecture. Scale bar, 500 µm. **(E)** Rat brain subcellular fractionation probed for MDGA1, postsynaptic excitatory marker PSD-95, and presynaptic marker synaptophysin. PSD: postsynaptic density. **(F)** Representative confocal images of dendritic segments from dissociated hippocampal neurons at different times in culture (7, 14, and 21 DIV) that were immunolabeled with MDGA1 antibody, and counterstained for either PSD-95 and VGLUT1. **(G, H)** Quantification of the co-localization level and area overlap between endogenous MDGA1 and the excitatory post-synaptic marker PSD-95, as a function of time in culture. Data represent mean ± SEM from three independent experiments, and were compared by a Kruskal–Wallis test followed by Dunn’s multiple comparison test (***P < 0,001). Scale bar, 10 µm.

We then examined the sub-cellular surface distribution of endogenous MDGA1 in dissociated rat hippocampal cultures at different developmental stages (DIV 7, 14, 21), by performing live staining of neurons with MDGA1 antiserum before fixation and counter immuno-labelling of the excitatory pre- and post-synaptic markers VGLUT1 and PSD-95, respectively (Figure 1F). Live labelling with this antibody was first tested in COS-7 cells expressing recombinant MDGA1 or MDGA2 constructs. Strong surface staining with the MDGA1 antiserum was observed in cells expressing MDGA1, but not in cells expressing MDGA2, validating this application and demonstrating no cross-reactivity of the antibody (Supplementary Fig. 2). In neurons, the MDGA1 staining revealed many sub-micron clusters, most likely a consequence of artifactual MDGA1 aggregation due to live incubation with the divalent polyclonal antibody ^21^. Those small puncta were distributed all over the dendritic shaft, and present but not particularly enriched at excitatory synapses. Quantitatively, the fraction of post-synaptic densities containing MDGA1 clusters was 45% and 40% at DIV 7 and 14, respectively, and decreased to 25% at DIV 21 (Figure 1G), suggesting that MDGA1 is leaving mature synapses and/or that the overall MDGA1 level decreases at later time points, as shown by RT-qPCR and Western blot analyses (Supplementary Fig. 1A-D). Among those synapses that contained MDGA1, the area overlap between PSD-95 and MDGA1 was around 20-30%, pointing to a minor occupancy of excitatory synapses, with no significant effect of neuronal maturation.

### Recombinant MDGA1 and MDGA2 are homogeneously distributed in dendrites at the nanoscale level

Next, we examined the nanoscale membrane organization of MDGAs using super-resolution microscopy. The formation of small endogenous MDGA1 aggregates observed upon live antibody labelling prevented a reliable estimation of MDGA distribution, as previously documented for NLGN1 ^21^. Moreover, we were lacking a good antibody to MDGA2 for surface staining. Thus, to monitor the precise localization of MDGAs expressed at near endogenous levels, we knocked-down native MDGA1 or MDGA2 with previously published shRNAs ^26, 31^ also validated in our conditions (Supplementary Fig. 3), and rescued them with recombinant rat MDGA1 or MDGA2 bearing the short N-terminal biotin acceptor peptide (AP), which is biotinylated upon the co-expression of biotin ligase (BirA^ER^) ^37^. We also included in the electroporation condition the excitatory post-synaptic marker Homer1c-DsRed. We then performed direct STochastic Optical Reconstruction Microscopy (dSTORM) experiments ^38^ after high density live labeling of AP-MDGA1/2 with monomeric streptavidin (mSA) ^39, 40^ conjugated to Alexa647 (100 nM concentration), reaching an optical resolution of about 30 nm. Since MDGAs are GPI-anchored membrane molecules, we electroporated neurons with GFP-GPI as a control, and labeled them with an anti-GFP nanobody also conjugated to Alexa647, a strategy previously validated for GFP-neurexin1β ^21^. Using this approach, AP-MDGA1 and AP-MDGA2 displayed a fairly uniform distribution at DIV 10 and 14, filling the whole dendritic shaft without specific accumulation at synapses, similarly to the negative control GFP-GPI (Figure 2A,C). In the post-synapse labeled by Homer1c-DsRed, MDGAs and GFP-GPI displayed a disperse localization (insets). For comparison, AP-NLGN1 expressed under similar replacement conditions (shRNA + rescue) and labeled identically with mSA-Alexa647, showed a stronger accumulation at synapses as previously shown ^21^. Synaptic enrichment at DIV 10 and 14 was around 1.3 and 1.5 for both MDGAs and GFP-GPI, and significantly higher for NLGN1 (2.3 and 2.7, respectively) (Figure 2B,D). These data show that MDGAs are not particularly enriched at excitatory synapses, and their differential localization with respect to NLGN1 suggest that the majority of NLGN1 molecules accumulated at post-synapses are not associated to MDGAs.

**Figure 2.**
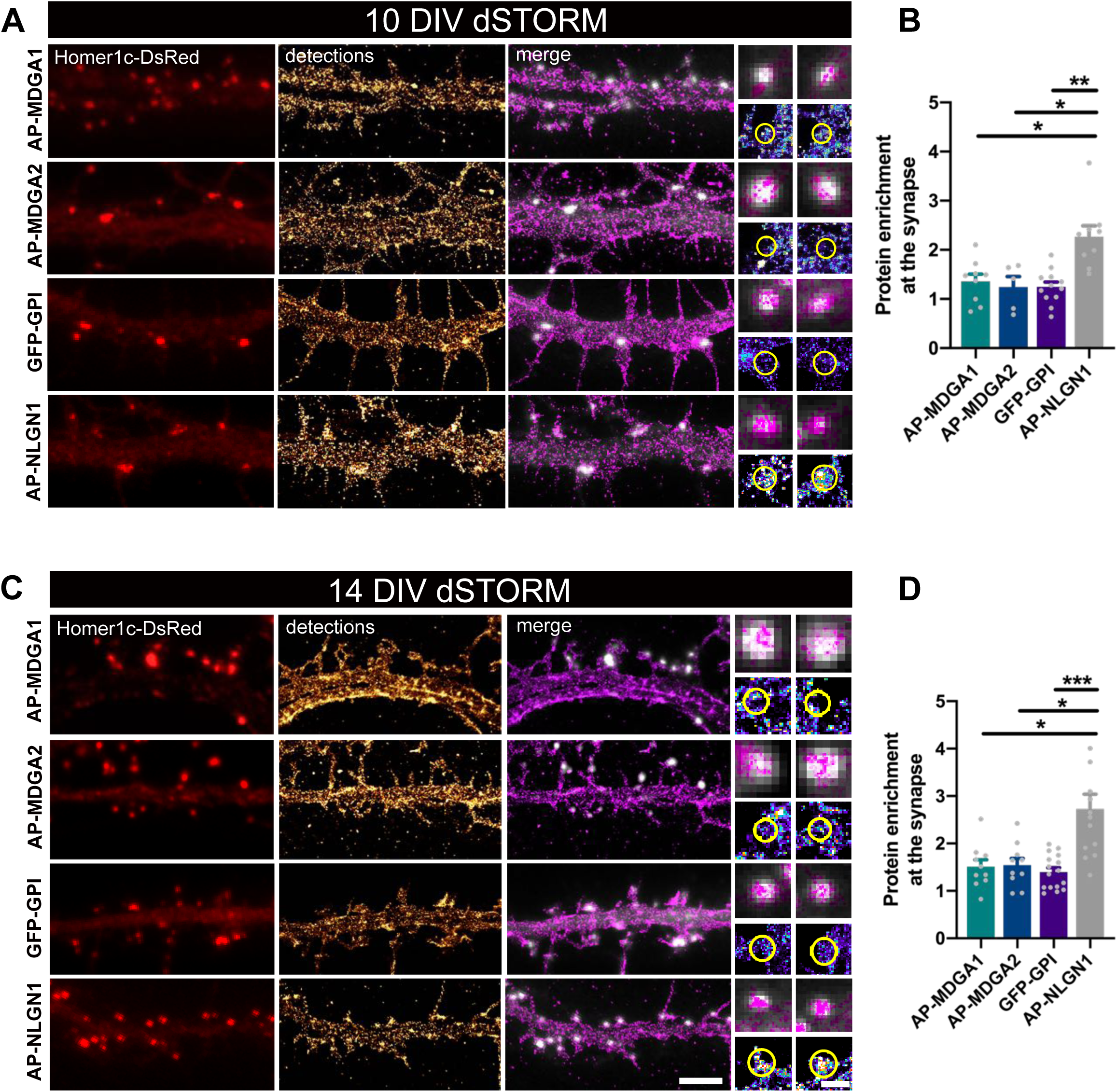
Nanoscale distribution of MDGA1 and MDGA2 in the neuronal membrane. Hippocampal neurons were electroporated at DIV 0 with a combination of shRNAs to MDGA1 or MDGA2, rescue AP-MDGA1 or AP-MDGA2 (respectively), biotin ligase BirA^ER^, and Homer1c-DsRed. Alternatively, neurons were electroporated with shRNA to NLGN1, rescue AP-NLGN1, biotin ligase BirA^ER^, and Homer1c-DsRed, or with GFP-GPI and Homer1c-DsRed. dSTORM experiments were performed at DIV 10 or 14, after labelling neurons with 100 nM Alexa647 conjugated mSA (for AP tagged MDGAs and NLGN1) or Alexa647 conjugated GFP nanobody (for GFP-GPI). **(A, C)** Representative images of dendritic segments showing Homer1c-DsRed positive synapses (in red), the super-resolved localization map of all AP-MDGA1, AP-MDGA2, GFP-GPI, or AP-NLGN1 single molecule detections (gold), and merged images (Homer1c-DsRed in white and detections in magenta). Scale bars 10µm. Insets on the right show zoomed images of different examples of Homer1c-DsRed positive puncta overlapped with localizations (magenta) or pseudo-coloured localizations in a synaptic area marked by a yellow circle. Scale Bars 1µm. **(B, D)** Bar plots representing the enrichment of AP-MDGA1, AP-MDGA2, GFP-GPI and AP-NLGN1 localizations at synapses. Values were obtained from at least three independent experiments and n > 5 for each experimental condition. Data were compared by a Kruskal–Wallis test followed by Dunn’s multiple comparison test (*P < 0.05; **P < 0.01; ***P < 0.001).

To rule out the possibility that the mSA probe was hindering the binding of MDGAs to NLGNs, and hence the penetration of MDGAs in synapses, we performed a series of control biochemical and immunocytochemical experiments. Streptavidin pull-down of proteins extracted from COS-7 cells expressing AP-MDGA1, BirA^ER^, and HA-NLGN1, followed by anti-MDGA1 and anti-NLGN1 immunoblots, revealed that AP-MDGA1 strongly recruits HA-NLGN1 (Supplementary Fig. 4B). This finding suggests that mSA, which is four times smaller than regular streptavidin ^39^, should easily access AP-tagged MDGAs bound to endogenous NLGN1 in neurons. Given the high sequence and structure similarity between MDGA1 and MDGA2, we expect AP-MDGA2 to also bind NLGN1 in this assay. To confirm that the interaction between MDGAs and NLGN1 also occurs when these molecules are bound to external probes in living cells, we performed cross-linking experiments using a mixture of a primary mouse anti-biotin and secondary anti-mouse antibodies in COS-7 cells expressing AP-MDGA1, HA-NLGN1 and BirA^ER^, or in cells expressing HA-MDGA2, AP-NLGN1 and BirA^ER^ (Supplementary Fig. 4C-F). In both cases, the fluorescent antibody clusters that aggregated AP-tagged proteins contained the HA-tagged co-expressed protein, demonstrating no hindrance caused by antibodies (which are much larger than mSA) on the MDGA-NLGN1 interaction. Strengthened by these controls, our dSTORM data strongly indicate that MDGAs are not enriched at post-synapses, supporting the concept that MDGAs do not bind NRXN-occupied NLGNs at synapses.

### Individual recombinant MDGA1 and MDGA2 are highly diffusive in the neuronal membrane

To characterize the surface dynamics of MDGAs at the individual level, we sparsely labelled biotinylated AP-MDGAs at the cell membrane using 1 nM mSA conjugated to the robust fluorophore STAR 635P, and performed single molecule tracking by universal Point Accumulation In Nanoscale Topography (uPAINT) (Figure 3), as described earlier ^21, 40^. Experiments were performed at DIV 8, 10, or 14, a time window of active excitatory synapse differentiation ^5^. As a control, we electroporated neurons with GFP-GPI and labeled them with an anti-GFP nanobody also conjugated to Atto 647N. At DIV 8, recombinant AP-MDGA1 and AP-MDGA2 diffused very fast in the dendritic membrane, showing a single peak of diffusion coefficient around 0.30 µm²/s, very similar to GFP-GPI (Figure 3A,B). Considering a small fraction around 20% of slowly mobile molecules, defined as molecules exploring an area smaller than the pointing accuracy of the optical system, i.e. D < 0.01 µm²/s ^21^, the median diffusion coefficient of the overall distribution was around 0.13-0.15 µm²/s across conditions (Figure 3G,H). Upon neuronal maturation (at DIV 10 and 14), the fraction of slowly mobile molecules increased for MDGA1 and MDGA2, with a concomitant decrease in median diffusion coefficient, while those parameters remained fairly constant for GFP-GPI (Figure 3C-H), suggesting a specific immobilization of MDGAs at these developmental stages.

**Figure 3.**
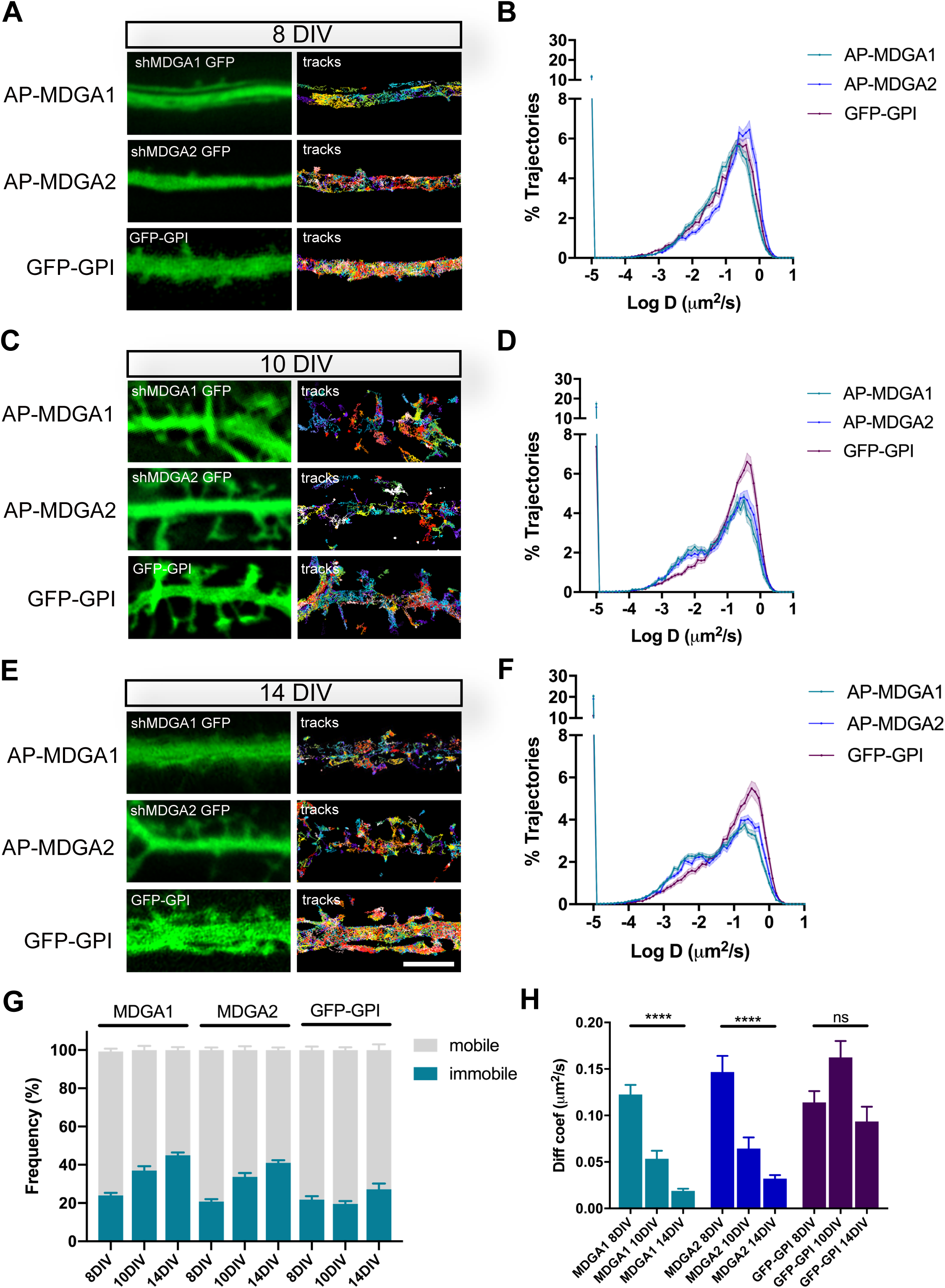
Lateral mobility of recombinant MDGAs across neuronal development. Dissociated rat hippocampal neurons were electroporated at DIV 0 with a combination of shRNAs to MDGA1 or MDGA2 (both carrying a GFP reporter), rescue AP-tagged MDGA1 or MDGA2 (respectively), and biotin ligase (BirA^ER^). Control neurons were electroporated with GFP-GPI. uPAINT experiments were performed at DIV 8, 10, or 14, after labelling neurons expressing AP-MDGA1 or AP-MDGA2 with 1 nM STAR 635P-conjugated mSA, and labelling neurons expressing GFP-GPI with 1 nM Atto 647N-conjugated anti-GFP nanobody. **(A, C, E)** Representative images of dendritic segments showing the GFP signal (green) and the corresponding single molecule trajectories (random colors) acquired during an 80 s stream, for the indicated time in culture. **(B, D, F)** Corresponding semi-log plots of the distributions of diffusion coefficients for AP-MDGA1, AP-MDGA2, and GFP-GPI, at the three different developmental times. **(G)** Graph of the mobile and immobile fractions of MDGA1, MDGA2, and GFP-GPI, as a function of time in culture. The threshold between mobile and immobile molecules was set at D = 0.01 µm²/s. **(H)** Graph of the median diffusion coefficient, averaged per cell, in the different conditions. Data represent mean ± SEM from at least three independent experiments (n > 10 for each experimental condition), and were compared by a Kruskal–Wallis test followed by Dunn’s multiple comparison test (**** P < 0.001).

### Individual recombinant MDGA1 and MDGA2 molecules are not trapped at synapses

Using the same set of data obtained from uPAINT experiments, we then examined the membrane domains explored by AP-tagged MDGA1 and MDGA2 in relation to the co-expressed post-synaptic marker Homer1c-DsRed, by constructing images integrating all single molecule localizations over time. We found that, at the individual level, neither AP-MDGA1 nor AP-MDGA2 molecules were particularly retained at synapses (Figure 4A,C), confirming the ensemble picture given by dSTORM. As shown in the insets, both MDGA1 and MDGA2 displayed a panel of localizations including: i) a complete absence from the post-synapse, ii) the formation of small clusters reflecting confined trajectories localized at the periphery of Homer1c-DsRed puncta, and iii) a more dispersed distribution filling the whole post-synapse (Figure 4A,C). For comparison, GFP-GPI exhibited essentially the third type of behavior, i.e. it explored the whole post-synapse with fast diffusion. Very rarely did MDGAs or GFP-GPI display confined trajectories at the core of the post-synaptic density like NLGN1 or LRRTM2 ^21^, suggesting an absence of synaptic retention. To directly compare the localization of MDGAs and LRRTM2, we expressed those molecules fused to an N-terminal V5 tag, and tracked them by uPAINT using a V5 Fab conjugated to STAR 635P. V5-MDGA1 and V5-MDGA2 showed similar peri- and extra-synaptic distribution as their AP-tagged counterparts (Supplementary Fig. 5), while V5-LRRTM2 exhibited striking post-synaptic confinement as previously reported ^21^. To quantitatively characterize the presence of individual AP-MDGA1 and AP-MDGA2 molecules at the post-synapse, we measured a parameter called synaptic coverage, and defined as the fraction of the area of Homer1c-DsRed puncta occupied by AP-MDGAs or GFP-GPI based on single molecule detections (Figure 4B,D). Synaptic coverage was only 20% at both 10 and 14 DIV, while it reached 40% for GFP-GPI, indicating that MDGAs dynamically explore only a minor fraction of the synaptic cleft.

**Figure 4.**
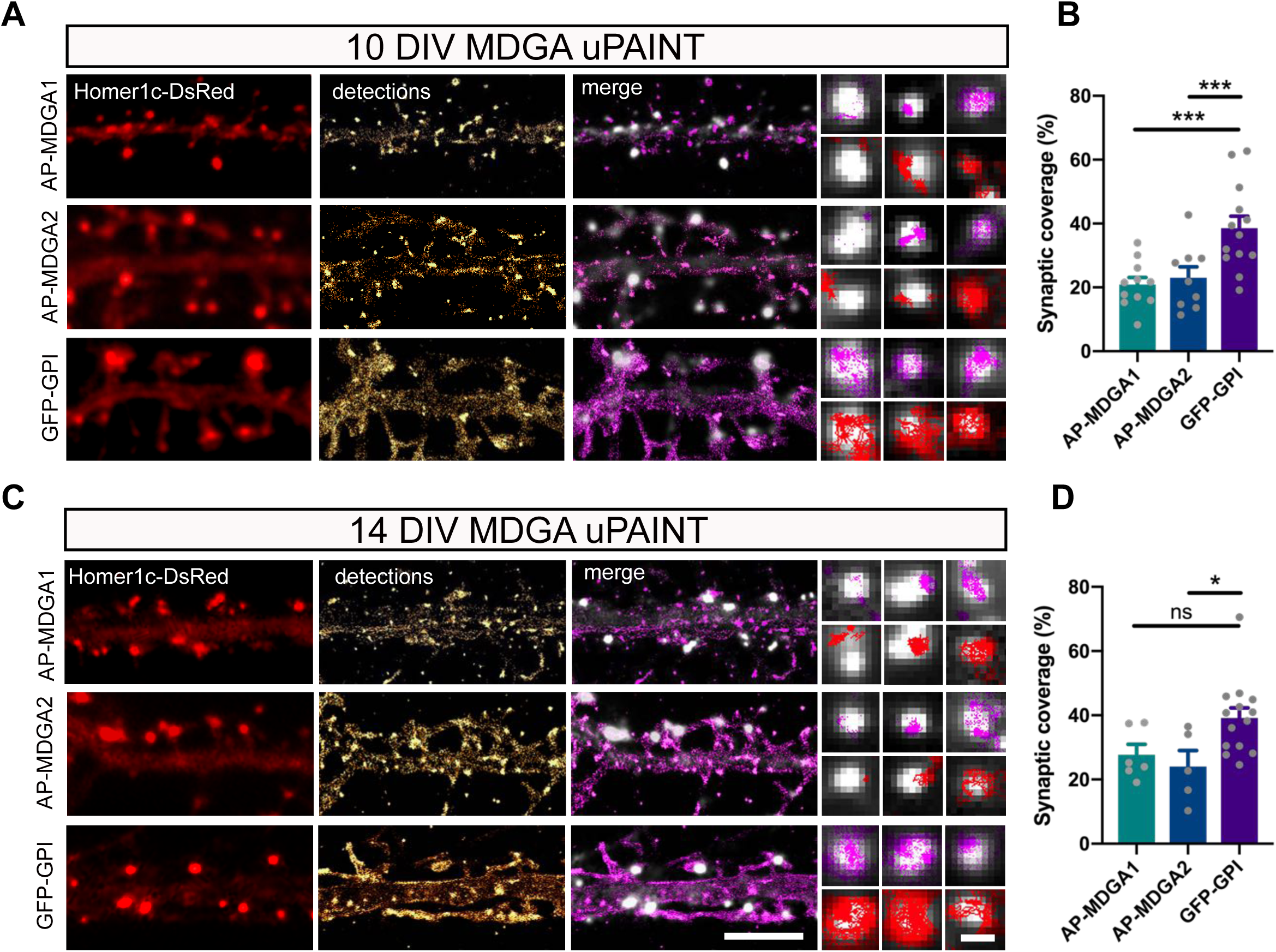
Single molecule localization of recombinant MDGAs with respect to post-synaptic densities. Hippocampal neurons were electroporated at DIV 0 with a combination of shRNAs to MDGA1 or MDGA2, rescue AP-MDGA1 or AP-MDGA2 (respectively), biotin ligase (BirA^ER^), and Homer1c-DsRed. Control neurons were electroporated with GFP-GPI and Homer1c-DsRed. uPAINT experiments were performed at DIV 10 or 14, after labelling neurons with 1 nM STAR 635P-conjugated mSA or Atto 647N-conjugated anti-GFP nanobody, respectively. **(A, C)** Representative images of dendritic segments showing the Homer1c-DsRed signal (red), the super-resolved localization map of all AP-MDGA1, AP-MDGA2, or GFP-GPI single molecule detections (gold), and the corresponding trajectories (magenta) super-imposed to Homer1c-DsRed (white). Scale Bars 10 µm. Insets represent zooms on individual post-synapses in the different conditions (Homer1c-DsRed in white, detections in magenta and trajectories in red). Scale bar 1 µm**. (B, D)** Bar plots representing synaptic coverage of AP-MDGA1, AP-MDGA2, or GFP-GPI at synapses, based on single molecule detections, for the two developmental stages (DIV 10 and 14), respectively. Values were obtained from at least three independent experiments and n > 10 for each experimental condition. Data were compared by a Kruskal– Wallis test followed by Dunn’s multiple comparison test (*P < 0.05, ***P < 0.001).

### MDGA2 knock-down increases synapse number and NLGN1 synaptic confinement

To characterize the influence of MDGAs on the behavior of their primary binding partner NLGN1, we knocked down MDGAs with shRNAs to MDGA1 (shMDGA1), to MDGA2 (shMDGA2), or to the non-related protein MORF4L1 as a control (shCTRL) ^26^. Neurons were co-electroporated at DIV 0 with these constructs together with Homer1c-DsRed. At DIV 10, a doubling in the density of Homer1c-DsRed puncta was observed in neurons expressing shMDGA2 relatively to shCTRL (Supplementary Fig. 6). No significant effect of shMDGA1 was observed on the density of excitatory post-synaptic clusters, confirming previous results ^26, 31^. For this reason, and considering the selective effects of MDGA2 on excitatory synapses reported earlier ^32^, we focused thereafter on the effects of MDGA2 on the dynamics, organization, and signaling mechanisms associated with NLGN1.

We first examined the diffusion properties of recombinant surface AP-NLGN1 sparsely labeled with STAR 635P-conjugated mSA with uPAINT. By comparing neurons at DIV 10 and 14, there was a shift in NLGN1 mobility towards lower diffusion coefficients, which reflects a synaptic immobilization of NLGN1 upon neuronal maturation, as previously reported ^21^. In DIV 10 neurons, shMDGA2 had no effect on the NLGN1 diffusion coefficient, whose distribution looked very similar to the shCTRL condition (Figure 5 **A-C)**. In contrast, at DIV 14, shMDGA2 decreased the global diffusion coefficient of NLGN1 as compared to shCTRL, in particular by reducing the mobile pool of NLGN1 molecules (the peak centered at D = 0.1 µm²/s), and concomitantly raising the fraction of confined NLGN1 molecules (peak at D = 0.01 µm²/s) that are most likely retained at synapses (Figure 5 **E-H)**. This effect was reversed upon the co-expression of an HA-MDGA2 construct resistant to shMDGA2. These data indicate that MDGA2 impairs the synaptic immobilization of NLGN1, i.e. MDGA2 knock-down exacerbates the confinement of NLGN1 that normally occurs during neuronal maturation.

**Figure 5.**
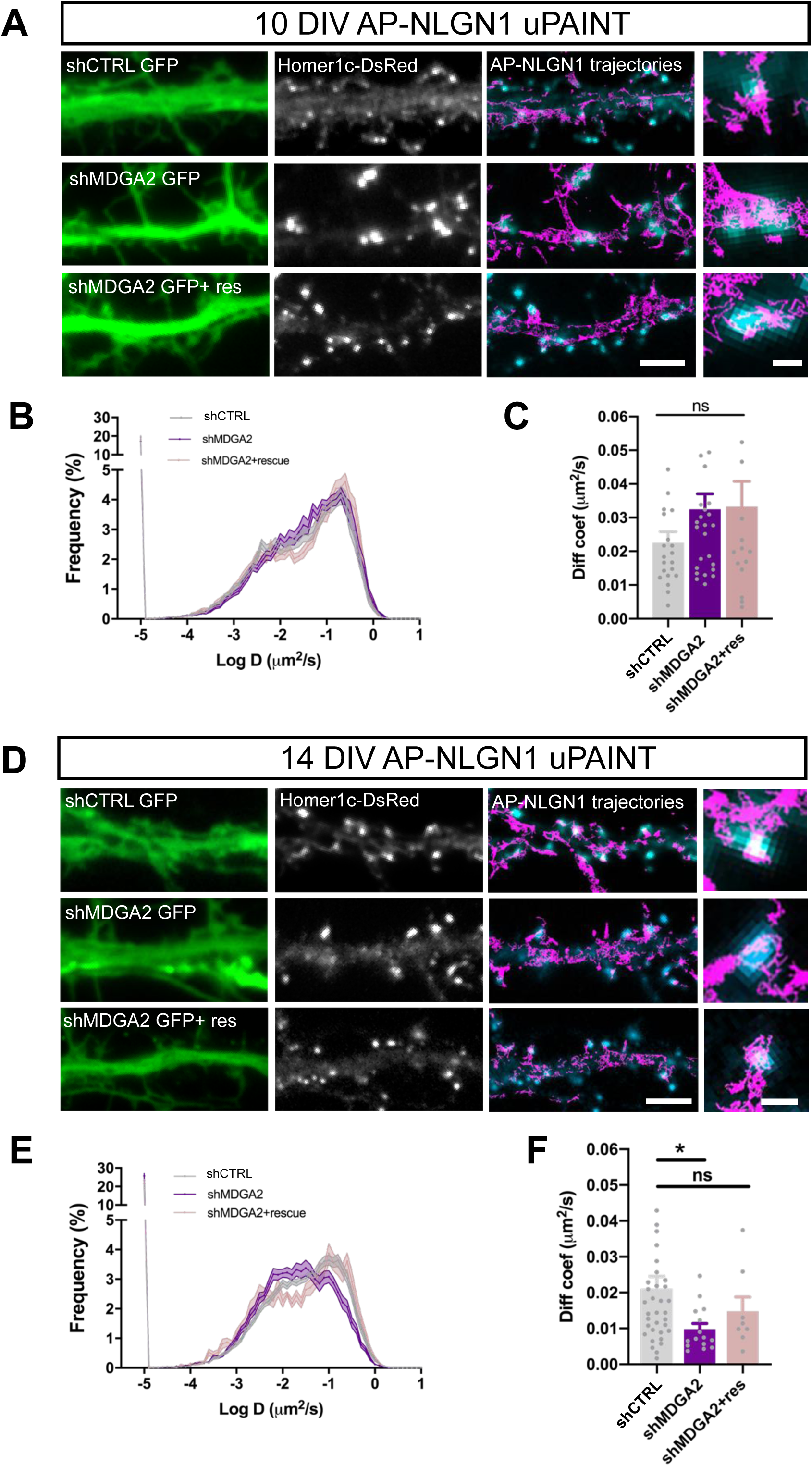
NLGN1 mobility and membrane localization upon MDGA2 knock-down. Neurons were electroporated at DIV 0 with AP-NLGN1, BirA^ER^ and Homer1c-DsRed, plus shCTRL, shMDGA2, or shMDGA2 + rescue HA-MDGA2, and imaged at DIV 10 or 14 using uPAINT. **(A, C)** AP-NLGN1 was sparsely labelled using 1 nM STAR 635P-conjugated mSA for single molecule tracking at 10 DIV (A) and 14 DIV (D). Super-imposed images of Homer1c-DsRed (cyan) and AP-NLGN1 trajectories (magenta) acquired during an 80 s stream are shown on the right of each panel. Scale bar, 5 μm. Insets represent zooms on individual post-synapses in the different conditions (Homer1c-DsRed in cyan, trajectories in magenta). Scale bar 1 µm**. (B, E)** Semi-log plot of the distribution of AP-NLGN1 diffusion coefficients. The curves represent the average of at least 15 neurons per condition, from three independent experiments. **(C, F)** Median diffusion coefficient of AP-NLGN1. Data represent mean ± SEM from at least 15 neurons per condition from three independent experiments, and were compared by a Kruskal–Wallis test followed by Dunn’s multiple comparison test (*P < 0.05).

Second, we examined the nanoscale distribution of surface AP-NLGN1 densely labeled with Alexa647-conjugated mSA using dSTORM. In DIV 10 neurons, there was no significant effect of shMDGA2 on NLGN1 enrichment at Homer1c-DsRed positive puncta compared to shCTRL (Figure 6A,B). In DIV 14 neurons, an increase from 3 to 4 in the synaptic enrichment of AP-NLGN1 was observed upon shMDGA2 expression as compared to shCTRL, albeit not significant (Figure 6C,D). This trend was reversed upon the co-expression of a MDGA2 construct resistant to the shRNA. Taken together, uPAINT and dSTORM data suggest that MDGA2 impairs the immobilization of NLGN1 at newly formed synapses, but not so much its intrinsic post-synaptic accumulation.

**Figure 6.**
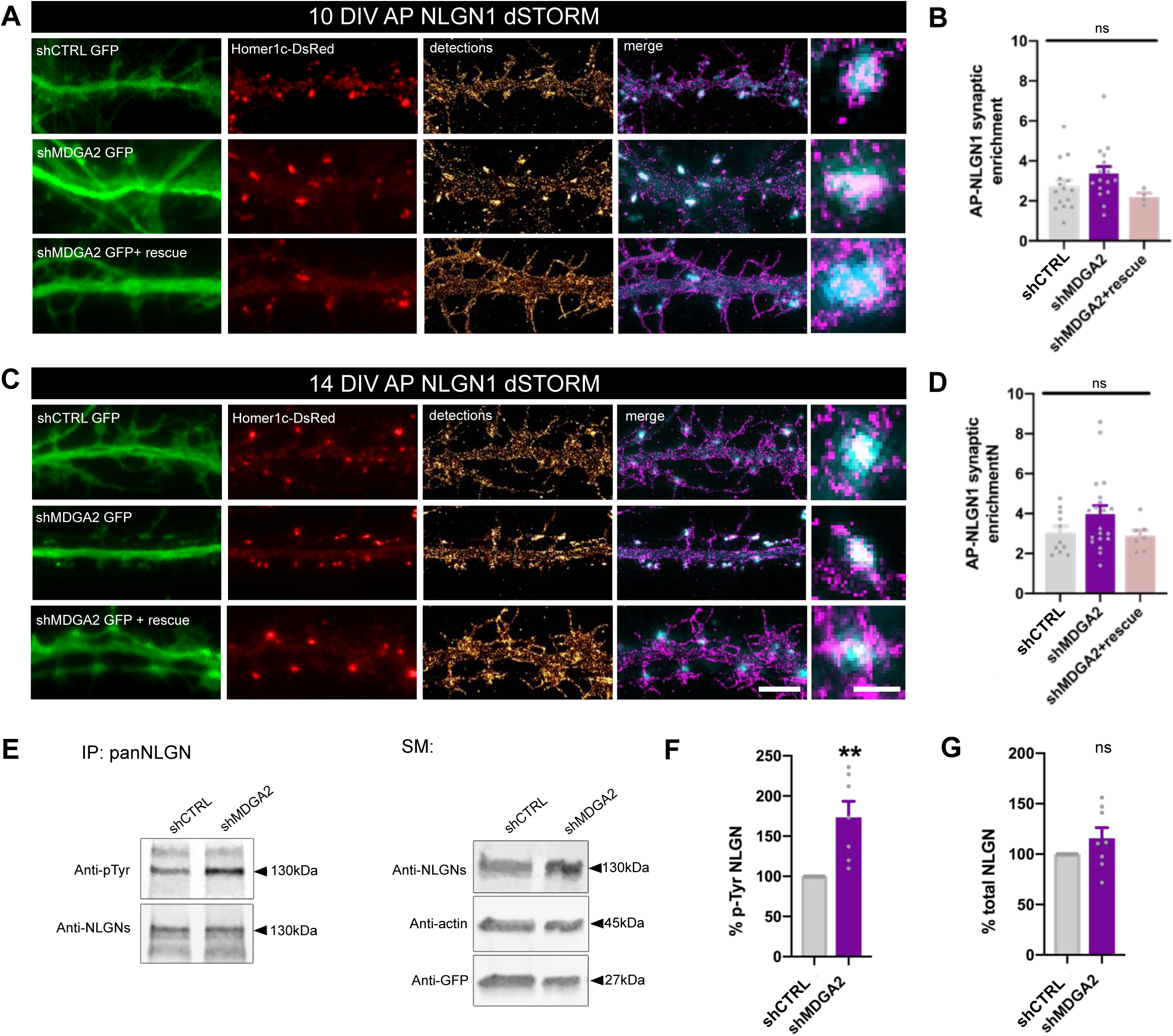
NLGN1 nanoscale membrane localization and phosphorylation upon MDGA2 knock-down. **(A, C)** Neurons were electroporated at DIV 0 with AP-NLGN1, BirA^ER^ and Homer1c-DsRed, plus shCTRL, shMDGA2, or shMDGA2 + rescue HA-MDGA2, and imaged at DIV 10 or 14 using dSTORM after densely labelling with 100 nM Alexa 647-conjugated mSA. Representative images of dendritic segments show the GFP reporter of shRNAs, Homer1c-DsRed (red) and the integration of all AP-NLGN1 single molecule localizations (gold). Merged images show Homer1c-DsRed (cyan) and AP-NLGN1 localizations (magenta). Scale bar, 10 μm. Insets on the right show zoomed examples of Homer1c-DsRed positive puncta overlapped with AP-NLGN1 localizations (magenta). Scale bar, 1μm. **(B, D)** Bar plots representing the enrichment of AP-NLGN1 at Homer1c-DsRed puncta. Data represent mean ± SEM from three independent experiments and were compared by a Kruskal–Wallis test followed by Dunn’s multiple comparison test (n > 4 at 10 DIV and n > 7 at 14 DIV for each construct). **(E)** Hippocampal neurons were electroporated at 0 DIV with shCTRL or shMDGA2 and cultured for 10 days. Protein extracts were immunoprecipitated with a pan NLGN antibody. Phosphotyrosine (pTyr) and total NLGN levels were detected by Western blot in the immunoprecipitation (IP) samples, and pan NLGN, actin and GFP were revealed in the starting material (SM). **(F)** Bar plots showing the average pTyrosine signal from the pan NLGN immunoprecipitate normalized to the total amount of immunoprecipitated NLGN. **(G)** Bar plots showing total amount of NLGNs in shCTRL and shMDGA2 electroporated cells. Data represent mean± SEM from 7 independent experiments and were compared by a t-test (**P > 0,01).

### MDGA knock-down enhances NLGN tyrosine phosphorylation

In view of our previous findings that the effects of NLGN1 on synapse number and AMPA receptor-mediated synaptic transmission are regulated by the phosphorylation of a unique intracellular tyrosine (Y782) in NLGN1 (Letellier et al., 2018;2020), we examined whether MDGAs could affect NLGN1 phosphotyrosine level. Our rationale was that by shielding NLGN1, MDGAs could impair the NLGN1 phosphorylation signaling mechanism which is dependent on NRXN binding (Giannone et al. 2013). We electroporated neurons at DIV 0 with shMDGA2 or shCTRL and analyzed the phosphotyrosine level of immunoprecipitated NLGNs by performing immunoblot at DIV 10, when NLGN phosphorylation is maximal ^3^. The NLGN phosphotyrosine level was almost two-fold higher in neurons expressing shMDGA2 compared to shCTRL, with no change in the total amount of NLGNs (Figure 6E-G). This result demonstrates that endogenous MDGAs negatively regulate NLGN tyrosine phosphorylation.

### MDGA2 knock-down reduces AMPA receptor diffusion

Given the previously reported effects of NLGN1 expression level and phosphotyrosine signaling on AMPA receptor surface trafficking and synaptic recruitment ^3, 18, 22, 34^, and seeing here the impact of MDGA2 knock-down on NLGN1 dynamics, localization, and phosphotyrosine level, we then questioned the role of MDGA2 on AMPA receptor surface diffusion. We electroporated hippocampal neurons at DIV 0 with shMDGA2 or shCTRL and tracked native AMPA receptors at the single molecule level by uPAINT upon sparse labeling with an antibody to the GluA2 N-terminal domain conjugated to Atto 647N ^22, 35, 41^. Expression of shMDGA2 significantly decreased the global AMPA receptor diffusion coefficient at DIV 10 compared to shCTRL (Figure 7A-D). Specifically, the mobile pool of AMPA receptors (D centered at 0.1 µm²/s) was reduced to the profit of slowly diffusing AMPA receptors (D < 0.01 µm²/s), most likely corresponding to synaptic receptors ^41^. This effect is consistent with the fact that shMDGA2 simultaneously increases the density of post-synapses (Supplementary Fig. 6), which act as trapping elements for surface-diffusing AMPA receptors ^34, 42^, resulting in an overall decrease in AMPA receptor mobility. At DIV 14, the distribution of AMPA receptor diffusion coefficients was shifted to the left as compared to DIV 10, reflecting AMPA receptor trapping at new synapses formed during this time interval (Figure 7E-H). Expression of shMDGA2 caused a further small decrease in diffusion coefficient, matching the observation that neurons expressing shMDGA2 show a trend for higher numbers of excitatory synapses at DIV 14 as compared to neurons expressing shCTRL (Supplementary Fig. 6C,D).

**Figure 7:**
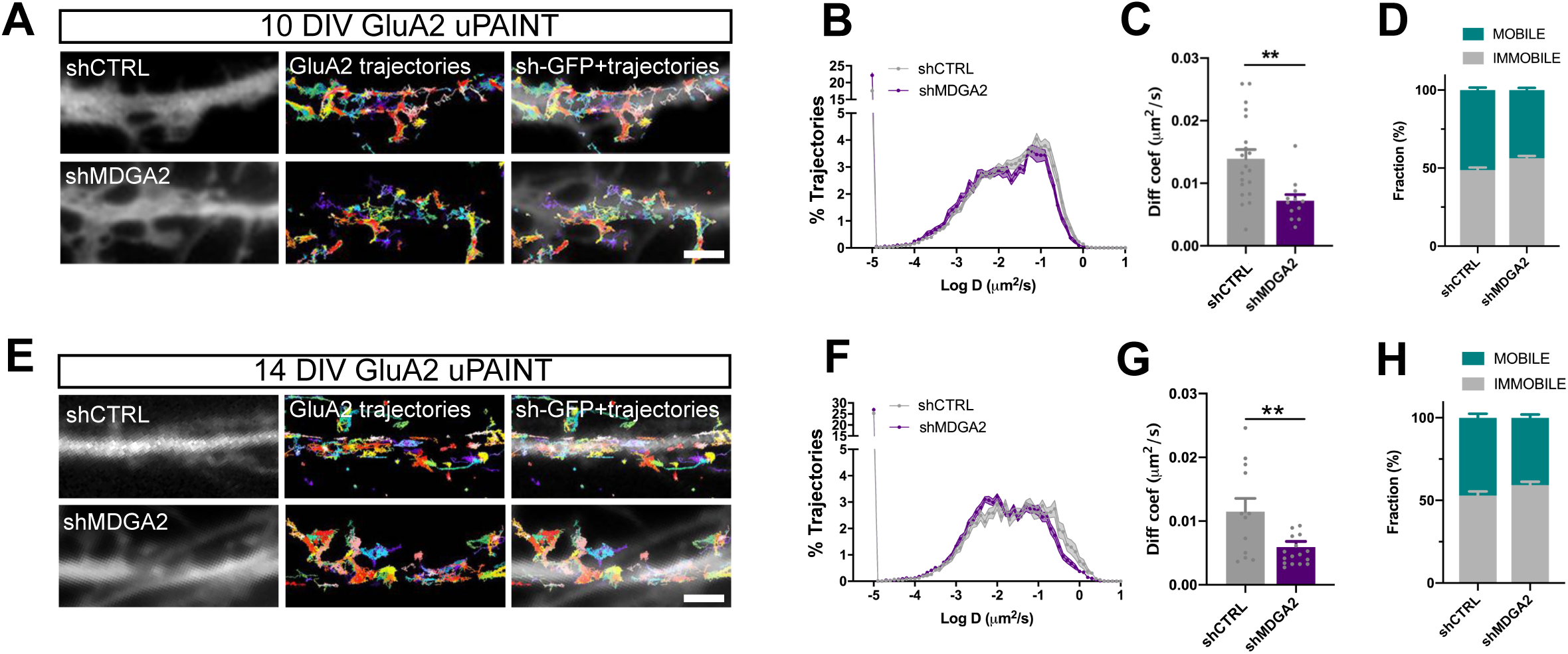
GluA2 membrane mobility upon MDGA2 knock-down. Neurons were electroporated at DIV 0 with shCTRL or shMDGA2-GFP, and imaged at DIV 10 or DIV 14 with uPAINT **(A, E)** GluA2 was sparsely labelled using a mouse monoclonal antibody anti GluA2 coupled to Atto 647N. GluA2 representative trajectories are shown multicolor. Scale bar, 2μm. **(B, F)** Semi-log plot of the distribution of GluA2 diffusion coefficients at 10 and 14 DIV, respectively. The curves represent the averages of at least 12 neurons per condition from three independent experiments. **(C, G)** Median diffusion coefficient of GluA2 at 10 and 14 DIV, respectively. Data represent mean ± SEM from at least 12 neurons per condition from three independent experiments, and were compared by a t-test (**P<0,01). **(D, H)** Bar plots of the immobile fraction of GluA2 in the three conditions, defined as the proportion of single molecules with diffusion coefficient D < 0.01 µm²/s.

### Both MDGA1 and MDGA2 knock-out promote excitatory post-synaptic maturation

Finally, to achieve a stronger suppression of MDGAs than that obtained with shRNAs and further highlight the roles played by MDGA1 and MDGA2 in excitatory synapse development, we designed new DNA vectors based on the CRISPR/Cas9 strategy to achieve single-cell knock-out of MDGA1 or MDGA2 in dissociated neurons ^43^. Specifically, hippocampal neurons were electroporated at DIV 0 with vectors containing the Cas9 gene, a guide RNA targeting either MDGA1, MDGA2, or a control sequence, plus a GFP or nuclear EBFP reporter. We first verified by genomic DNA cleavage that CRISPR was cutting the expected region of MDGA1 or MDGA2 genes only when the respective gRNA was present (Supplementary Fig. 7A). Moreover, after 10 DIV, we observed an 80% reduction of endogenous MDGA1 immunostaining in neurons expressing CRISPR-Cas9 and gRNA to MDGA1, compared to neurons expressing control gRNA, revealing MDGA1 knock-out efficiency (Supplementary Fig. 7B,C). We then evaluated the effects of MDGA1/2 knock-out on the number and area of individual excitatory pre- and post-synaptic areas immunolabeled for VGLUT1 and PSD-95, respectively. At DIV 10, an almost doubling in the number of PSD-95 puncta per unit dendrite length without a change in PSD-95 area, was observed in neurons expressing gRNAs to MDGA1 or MDGA2 relatively to control gRNA (Figure 8A-C). In the same conditions, only gRNA to MDGA2 caused a significant increase in the density of VGLUT1 puncta, and no change in area (Figure 8A, D, E). Those effects of gRNA to MDGA2 on both PSD-95 and VGLUT1 cluster density were abolished by the co-expression of a rescue MDGA2 vector, demonstrating the specificity of the mechanism. At DIV 14, no significant effects of gRNAs to MDGA1 or MDGA2, relative to control gRNA were found on PSD-95 or VGLUT1 cluster density or intensity (Supplementary Fig. 8). Together, these data show that both MDGA1 and MDGA2 mainly impair post-synaptic assembly in the early phase of synaptogenesis.

**Figure 8.**
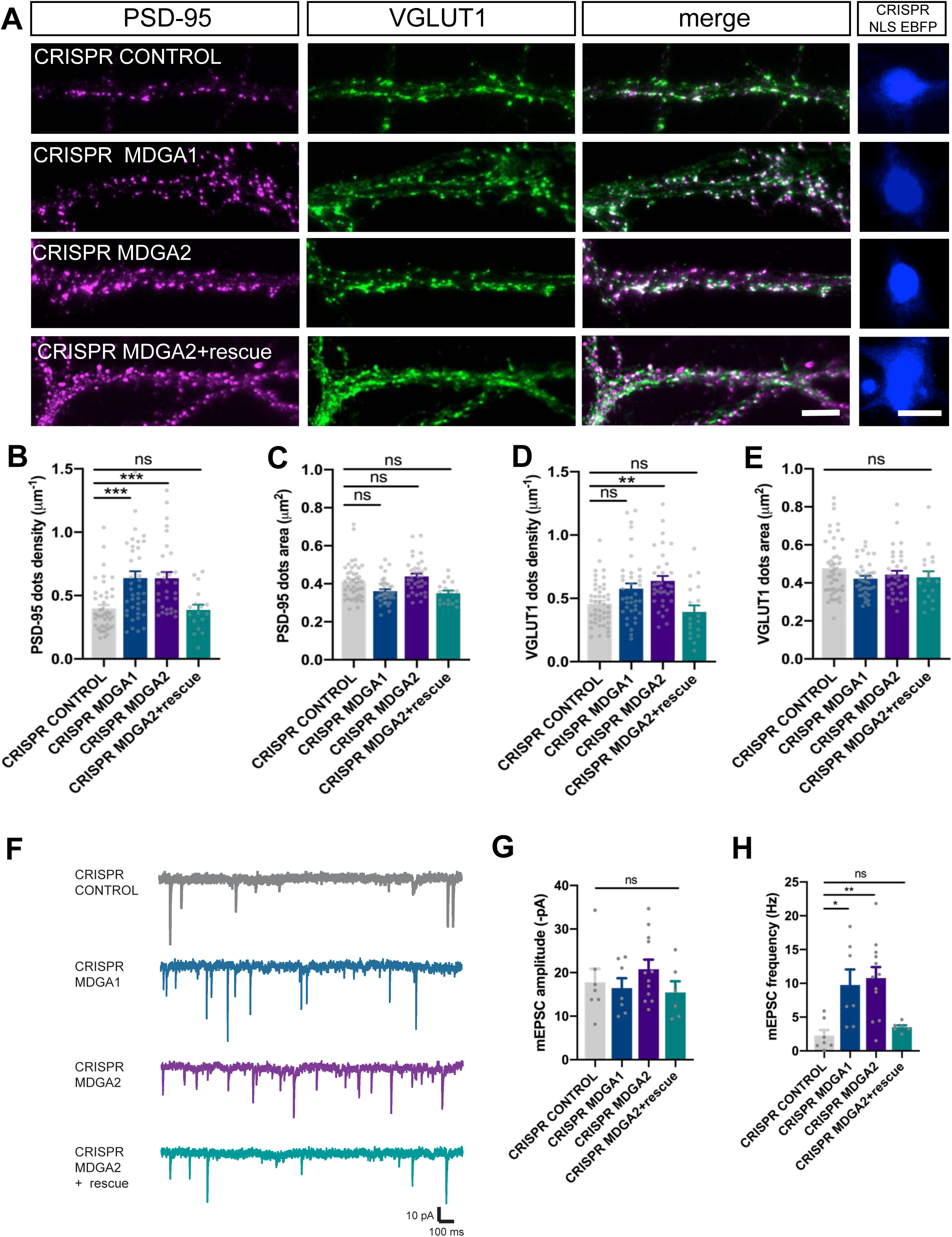
Effect of MDGA knock-out on synaptic density and transmission. Dissociated neurons where electroporated at 0 DIV with CRISPR/Cas9 control, CRISPR/Cas9 MDGA1, CRISPR/Cas9 MDGA2, or CRISPR/Cas9 MDGA2 plus HA-MDGA2 rescue. 10 DIV after plating, cultures were fixed, permeabilized, and endogenous PSD-95 and VGLUT1 were immunostained. **(A)** Representative images of dendritic segments showing PSD-95 staining (magenta), VGLUT1 staining (green), the merged images, and the nuclear EBFP control of CRISPR/Cas9 construct expression (blue), in the different conditions. Scale bars, 10 µm. **(B-E)** Bar plots showing the density per unit dendrite length and surface area of individual PSD-95 and VGLUT1 puncta, respectively, in the various conditions. Data represent mean ± SEM from at least three independent experiments, and were compared by a Kruskal–Wallis test followed by Dunn’s multiple comparison test (**P < 0.01; ***P < 0.001). **(F)** Representative traces of AMPA receptor-mediated mEPSC recordings from DIV 10 neurons expressing CRISPR/Cas9 control, CRISPR/Cas9 MDGA1, CRISPR/Cas9 MDGA2, or CRISPR/Cas9 MDGA2 plus HA-MDGA2 rescue, clamped at −70 mV in the presence of tetrodotoxin and bicuculline. **(G, H)** Bar graphs of mEPSC amplitude and frequency respectively, for each condition. Plots represent mean ± SEM from five independent experiments (each point represent one cell), and were compared by a Kruskal–Wallis test followed by Dunn’s multiple comparison test (*P < 0.05; **P < 0.01).

To examine the functional consequences of MDGA1 and MDGA2 knock-out on synaptic assembly, we measured AMPA receptor-mediated miniature EPSCs (mEPSCs) by performing whole cell patch-clamp recordings in neurons electroporated with either gRNAs to MDGA1 or MDGA2, or control gRNA. Neurons expressing gRNAs to MDGA1 or MDGA2 showed a three-fold increase in the frequency of AMPA receptor-mediated mEPSCs compared with neurons expressing control gRNA, while the combined expression of gRNA to MDGA2 and rescue MDGA2 abolished this effect (Figure 8F,H). No significant change in the amplitude of AMPA receptor-mediated mEPSCs was observed across conditions (Figure 8G). In parallel, endogenous surface AMPA receptors were live labeled with antibodies to the N-terminal of GluA1 subunits. There was no significant difference in GluA1 or GluA2 synaptic enrichment in neurons expressing CRISPR-Cas9 and gRNAs to MDGA1 or MDGA2, compared to neurons expressing CRISPR-Cas9 and control gRNA, despite an increase in the density of post-synaptic puncta as labeled by an intrabody to PSD-95, Xph20 ^44^ (Supplementary Fig. 9A-F). Together, these data suggest that knocking out either MDGA1 or MDGA2 affects the density of AMPA receptor-containing synapses, but not the actual amount of AMPA receptors per synapse.

## Discussion

In this study, we characterized the membrane localization of MDGAs and their role on the dynamics and signaling of their direct binding partner, NLGN1, as well as associated effects on synaptic differentiation and the recruitment of AMPA receptors. We demonstrate that MDGA1 and MDGA2 are essentially non-synaptically enriched molecules that exhibit fast diffusion in the dendritic membrane. Moreover, the knock-down of MDGAs increases synapse density and as a consequence reduces the surface mobility of both NLGN1 and AMPA receptors, increases AMPA receptor-mediated mEPSC frequency, and NLGN1 phosphosignalling. Thus, by shielding a fraction of NLGN1 from binding to pre-synaptic NRXNs, MDGAs negatively regulates NLGN function in excitatory synaptic differentiation (Figure 9).

**Figure 9.**
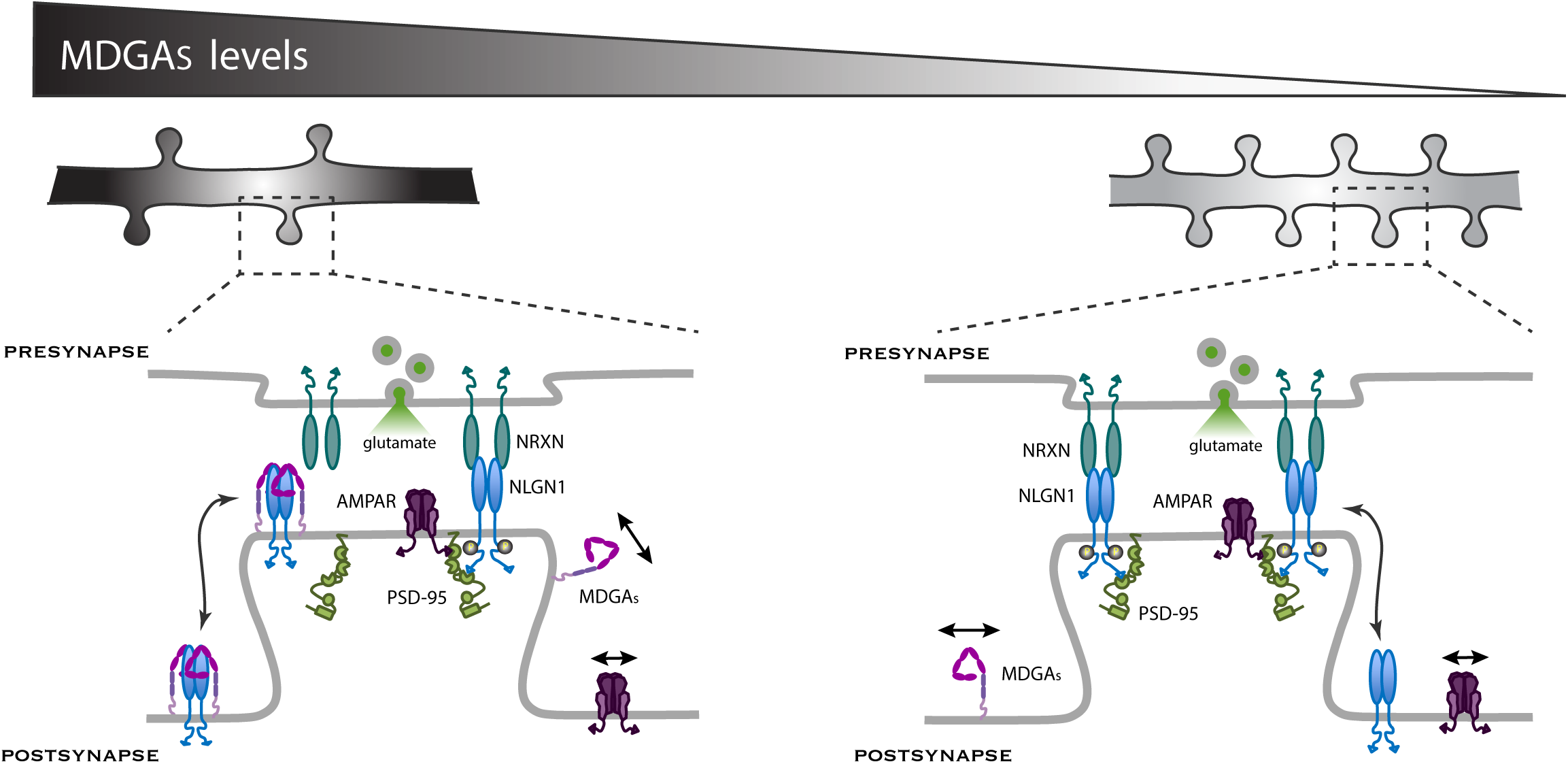
Working model for the role of MDGAs in synaptogenesis. Since NRXNs and MDGAs compete for the binding to NLGN1, the MDGA concentration acts as a key regulator of the signaling events downstream of the NRXN-NLGN1 interaction. When the MDGA level is low (in response to KD or KO), the preferential interaction of NLGN1 with NRXN favors NLGN1 tyrosine phosphorylation and the associated development of excitatory synapses containing AMPA receptors (right). When the MDGA level is high, the NRXN-NLGN1 interaction is weakened and the formation of excitatory synapses is delayed (left). MDGAs primarily regulate the overall density of NLGN1 and AMPA receptor modules, but not the actual amount of these molecules at individual synapses.

We first examined the localization of MDGAs in hippocampal tissue and dissociated cultures. The generation of specific antibodies allowed us to examine the distribution of native MDGA1, which showed strong expression levels in the neuropil of the CA region of the hippocampus, confirming previously shown results of *in situ* hybridization of MDGA1 mRNAs and staining of β-galactosidase activity expressed from the *Mdga1* locus ^27, 30, 32^. MDGA1 immunostaining in dissociated cultures at DIV 14 showed that a large fraction of excitatory synapses did not contain MDGA1. Recombinant MDGA1 and MDGA2 expressed by rescuing endogenous MDGA levels also showed no preferential retention at Homer1c-positive puncta, in contrast with the positive controls NLGN1 and LRRMT2 labeled similarly and showing strong post-synaptic accumulation, as previously described ^21^. Previous studies also reported a small colocalization extent of either YFP-MDGA1 or HRP-MDGA2 with excitatory synaptic markers ^26, 31^. Our results show that individual MDGAs often localized as transient sub-micron clusters at the periphery of Homer1c puncta, representing the confinement area of a small number of labeled molecules. The nature of these domains is unclear, but might represent a transition zone where NLGN1 could switch between MDGA-bound and NRXN-bound states, with the potential existence of mixed NLGN1 dimers that would exhibit specific mobility properties. In any case, the sum of this MDGA-rich peri-synaptic compartment plus the diffusive pool of MDGAs in synapses might represent enough material to be detected in synaptosomes (this study) and in the synaptic cleft proteome ^31^. When sampled at saturating labeling density using dSTORM, MDGAs showed a rather diffuse localization in the shaft and at synapses both at DIV 10 and 14, with modest synaptic enrichment as the negative control GFP-GPI. As compared to uPAINT, which gives information on the membrane dynamics of a subset of sparsely labeled molecules, dSTORM performed after saturating live labeling provides a snapshot of the whole population of MDGAs that integrates over time many transient confinement areas into a single 2D projection image. Thus the localization of molecules observed in dSTORM seems naturally more homogeneous, as previously reported for NLGN1 ^21^.

In addition, AP-MDGA1 and AP-MDGA2 exhibited fast membrane diffusion throughout dendrites, even within post-synaptic sites, indicating no particular molecular retention at synapses. These observations are compatible with the fact that MDGAs lack an intracellular C-terminal domain such as those present in NLGN1 or AMPA receptor auxiliary subunits, which bind PDZ domain-containing scaffolding proteins that stabilize them at synapses ^34, 45, 46^. A mild confinement of MDGAs outside synapses was observed upon neuronal maturation, which might be due to the fact that a fraction of MDGAs bind to extra-synaptic NLGN clusters ^47^ or to another unknown protein, e.g. through their Ig4-6 domains ^27^. These domains might also be due to interactions of the GPI anchor of MDGAs to the lipid microenvironment present in some membrane microdomains such as lipid rafts ^48, 49^. By comparing side by side the distributions of diffusion coefficients for MDGAs and NLGN1, we estimate that around 12% of the highly mobile extra-synaptic MDGAs are not bound to NLGN1, otherwise MDGAs would naturally adopt the slower diffusion of NLGN1. Furthermore, the fact that MDGAs do not accumulate at synapses like NLGN1 over neuronal development (from DIV 10 to 14) suggests that synaptic NLGN1 is not bound to MDGAs, but instead to NRXNs which display high local concentration at pre-synapses ^21, 50^. Thus, by preventing dendritic NLGN1 from aberrantly binding to the fraction of freely-diffusing NRXNs at the surface of contacting axons ^21, 51^, native MDGAs seem to protect neurons from forming synapses too quickly.

Using either previously reported shRNAs or newly generated CRIPSR/Cas9 constructs, we showed that MDGA2 delays excitatory synaptic development and AMPA receptor-dependent transmission, consistent with the *in vivo* KO of *Mdga2* ^32^. These effects were accompanied by a global decrease of NLGN1 diffusion and some increased confinement at post-synapses, supporting the idea that by losing its MDGA partner, NLGN1 is more available to bind pre-synaptic NRXNs and thereby accelerates synapse formation. Indeed, the effects of MDGA2 KD and KO were more prominent at DIV 10 during the active phase of synaptogenesis, and barely detectable at DIV 14. This result agrees with the lack of effect of shMDGA2 previously seen on excitatory synapse density in DIV 15 neurons ^31^. We found more contrasted results with MDGA1, i.e. the CRISPR/Cas9 strategy significantly increased excitatory post-synaptic density and AMPA receptor-mediated mEPSC frequency at DIV 10 (but not VGlut1 puncta density), while shRNA to MDGA1 had little effect at both DIV 10 and DIV 14, in agreement with previous reports ^26^. This discrepancy might be due to the fact that CRISPR/Cas9 suppresses MDGA1 expression more strongly than shMDGA1. Indeed, in previous studies, effects of MDGAs on excitatory and inhibitory synapse development were seen only when both MDGA1 and MDGA2 were knocked down ^27, 31^, suggesting that the overall level of MDGAs is an important parameter in these experiments. The reported increase in inhibitory - but not excitatory - synapses in hippocampal CA1 neurons of *Mdga* KO mice ^30^ might be due to circuit effects, and differences might arise when analyzing different cell types or different brain regions, considering that the relative expression levels of MDGA1 and MDGA2 may vary within neuronal types. In any case, crystal structures and affinity measurements support the concept of a stable MDGA1/NLGN1 complex ^25, 28, 29^, which should be compatible with the fact that endogenous MDGA1 can interact with NLGN1 as strongly as MDGA2 to impair excitatory synapse formation.

We also demonstrated a selective increase in the phosphotyrosine level of NLGNs upon MDGA knock-down. This observation relates to our previous findings that NLGN1 can be phosphorylated at a unique intracellular tyrosine residue (Y782), and that the NLGN1 phosphotyrosine level regulates the assembly of excitatory post-synaptic scaffolds in a NRXN-dependent fashion ^16^. Moreover, using either the expression of NLGN1 point mutants or the optogenetic stimulation of endogenous NLGN1 phosphorylation, we recently showed that a high NLGN1 phosphotyrosine level is associated with the selective increase in excitatory synapse number and AMPA receptor-mediated synaptic transmission ^3, 18^. Thus, we propose the model that by occupying NLGN1, MDGAs inhibit the NRXN-induced phosphotyrosine signaling pathway associated with NLGN1 and thereby delay the assembly of functional excitatory synapses. Since NLGN3 can also be tyrosine phosphorylated in neurons ^3^, a contribution of NLGN3 to the increase in phosphotyrosine level seen upon MDGA2 KD is possible given that we used a pan NLGN antibody to reach efficient immunoprecipitation. However, the fact that MDGA binds 10-fold more weakly to NLGN3 than to NLGN1 *in vitro* ^25^, and that the effects of NLGN tyrosine phosphorylation on post-synaptic differentiation are not seen in cultures from *NLGN1* KO mice ^18^ suggest that NLGN3 might only play a minor role in this process.

The decrease in global AMPA receptor diffusion observed upon MDGA2 knock-down can be linked to the parallel increase in the density of post-synaptic areas. Indeed, a similar decrease in AMPA receptor diffusion was seen across neuronal development in culture or upon the over-expression of NLGN1, which both enhance the number of post-synapses that act as trapping elements for surface diffusing AMPA receptors ^34, 42^. Upon MDGA2 knock-down, a transient immobilization of surface-diffusing AMPA receptors is expected to occur at newly formed synapses enriched in NLGN1, resembling what was previously observed at micro-patterned dots coated with NRXN1β-Fc ^35^. Interestingly, the actual content of AMPA receptors per post-synapse did not seem to be modified by MDGA knock-out, since both the synaptic AMPA receptor enrichment and the amplitude of AMPA receptor-mediated EPSCs remained similar to control conditions. However, the density of synaptic puncta as well as the frequency of AMPA receptor-mediated mEPSCs were significantly enhanced by MDGA knock-out, suggesting that the new synapses that had appeared contained functional AMPA receptors. This situation is quite similar to NLGN1 over-expression that increases the number of synapses and the frequency, but not the amplitude, of AMPA receptor-mediated mEPSCs ^3, 52^. In both cases (MDGA knock-down or NLGN overexpression), AMPA receptors seem to be inserted in novel synapses as individual units or modules, most likely influenced by the presence of _NLGN1_ ^22, 41, 53, 54^.

Given the strong effects caused by MDGA loss-of-function on synaptic differentiation, the next challenge would be to determine which biological processes regulate endogenous MDGA levels in specific neuron types across development. One interesting factor might be constitutive neuronal activity that can influence synaptic protein expression levels, e.g. by modulating microRNAs ^55, 56^. Indeed, MDGA1 transcripts were recently found to be upregulated in response to chronic synaptic activity blockade ^57^. Moreover, the action of MDGAs might be finely tuned by other proteins associated to the NRXN-NLGN trans-synaptic complex, including hevin and SPARC that are secreted by astrocytes ^58^. Interestingly, the presence of NLGNs in astrocytes offers an additional level of regulation of synapse development through such a network of proteins ^59^. Finally, genetic mutations identified in patients with autism and leading to alterations in MDGA levels ^60^, are expected to cause profound changes in synapse differentiation such as the ones shown here.

## Acknowledgements

We thank AM. Craig (University of British Columbia, Vancouver), A. Ting (Stanford University, Palo Alto), and P. Scheiffele (Biozentrum, Basel) for the generous gift of plasmids, T. Yamamoto (Kagawa University, Japan) for providing *Mdga1* KO mice lines, E. Gouaux (OSHU, Vollum Institute, Portland) for the gift of anti-GluA2 antibody, S. Benquet and M. Munier in the team for molecular biology, R. Sterling and J. Girard for logistics, the Cell Biology Core facility of the Institute (C. Breillat, N. Retailleau, E. Verdier, N. Chevrier), C. Lemoigne for probe production, J.B. Sibarita, A. Kechkar and C. Butler (IINS) for the generous gift of the PALM-Tracer and WAVE-Tracer single molecule detection programs, M. Mondin and C. Poujol (Bordeaux Imaging Center) for providing image analysis macros, AM. Craig, J. Elegheert (IINS) and N. Brose (Max Planck Institute, Goettingen) for scientific discussions. Confocal microscopy was done at the Bordeaux Imaging Center, part of the FranceBioImaging national infrastructure (ANR-10-INBS-04-0). The protein quantitation and western blot analysis were done in the Biochemistry and Biophysics Core Facility of the Bordeaux Neurocampus (BioProt) with the help of Yann Rufin.

This work received funding from the Fondation pour la Recherche Médicale (“Equipe FRM” DEQ20160334916), French Ministry of Research, Agence Nationale de la Recherche (grants « SynSpe » ANR-13-PDOC-0012-01, and « SyntheSyn » ANR-17-CE16-0028-01), ERA-NET Neuron “Synpathy” (ANR-15-NEUR-0007-04), Investissements d’Avenir Labex BRAIN (« SynOptoGenesis » ANR-10-LABX-43 ERC grant DynSynMem (787340) and grants from the conseil Régional de nouvelle Aquitaine. The BioProt facility is funded by the Labex BRAIN (ANR-10-LABX-43).

## Materials and Methods

### DNA constructs

Rat V5-MDGA1, V5-MDGA2, HA-MDGA1, HA-MDGA2, shMDGA1, shMORB (shCTRL), sh-RNA resistant HA-MDGA1 (rescue) constructs as described previously ^25, 26^ were kind gifts from A.M. Craig (University of British Columbia, Vancouver, OR). Mouse AP-tagged NLGN1, biotin ligase (BirA^ER^), shMDGA2, and mApple-V5-MDGA2 rescue ^31^ were gifts from A. Ting (Stanford University, CA). AP-MDGA1 and AP-MDGA2 were generated by replacing the V5 tag of the V5-MDGA1 and V5-MDGA2 constructs respectively by the 14 amino acids AP tag (5’GGCCTGAACGAtATCTTCGAGGCCCAG AAGATCGAGTGGCACGAG3’) using the HD-In-Fusion kit (Takara). The linker 5’GGAGGATCAGGAGGATCA3’ was added after the AP tag. AP-MDGA1 and AP-MDGA2 rescue constructs were generated by inserting the mutations responsible for the resistance to the respective shRNAs obtained from HA-MDGA1 and mApple-V5-MDGA2 rescue constructs, respectively, using the HD-In-Fusion kit. HA-MDGA2 rescue was created by replacing the AP tag from the AP-MGDA2 rescue construct by the HA tag using the HD-In-Fusion kit.

The CRISPR target sequences were all 20-nucleotide long and followed by a protospacer adjacent motif (PAM). The first step in the design of gRNAs involved identification of the best sequence to target. For MDGA1, we chose the more efficient gRNA proposed by the online software ChopChop (https://chopchop.cbu.uib.no/). For MDGA2, we chose to target the exon1, near ATG ^61^. The guide RNA (gRNA) sequence for MDGA1 was 5’ CTTCAACGTACGAGCCCGGG 3’, and for MDGA2 5’ TCACTAAACAGCTCCCCCGA 3’. As a control we used a sequence from a gecko bank, sequence: 5’ ATATTTCGGCAGTTGCAGCA 3’. gRNAs were cloned into the vector pSpCas9(BB)-2A-GFP (PX458) (Addgene cat#48138). The CRISPR MDGA2 resitant sequence was the same as for shMDGA2 since the gRNA for MDGA2 was directed to the signal peptide, which is absent in the HA-MDGA2 rescue decribed in the previous paragraph. Short hairpin RNA to murine NLGN1 (shNLGN1) ^62^ and HA-NLGN1 were gifts from P. Scheiffele (Biozentrum, Basel). shRNA-resistant AP-tagged NLGN1 (AP-NLGN1res) was described previously ^3, 21^. Homer1c-DsRed was described previously ^34^. The GFP-GPI construct was previously described ^48^. Xph20-GFP and Xph20-mRuby2 (Addgene#135530 pCAG_Xph20-eGFP-CCR5TC, #135531 pCAG_Xph20-mRuby2-CCR5TC) have previously been described ^63^.

### Production and fluorophore conjugation of probes

The anti-GFP nanobody and mSA were produced as previsouly described ^21, 40^. Briefly, the two proteins were expressed in E. Coli by auto-induction at 16 °C. Both proteins were purified by affinity chromatography using their polyhistidine tags in native and denaturing conditions for the nanobody and mSA respectively. After dialysis in PBS and concentration to ∼1 mg.mL^-1^, the proteins were coupled with 3-6 equivalents of the dyes in their activated ester form. Dyes used were Atto647N (Atto-Tec), Star635P (Abberior) and Alexa Fluor 647 (ThermoFisher). Excess unreacted dye was removed using a desalting column and the dye-conjugated probes were further purified to homogeneity by size-exclusion chromatography. Probes were concentrated and flash-frozen for storage at −80 °C until use. The anti-GluA2 antibody, clone 15F1 (gift from Eric Gouaux, OSHU, Vollum Institute, Portland), and the V5 tag recombinant Fab fragment (Abnova, RAB00032) were coupled NHS-derived dyes using the same protocol as above but without the size-exclusion chromatography purification step.

### MDGA1 recombinant protein production and rabbit polyclonal antiserum

For antibody production, mouse MDGA1 cDNA lacking signal peptide, GPI anchor site, and propeptide (amino acids 19-932; Uniprot ID# Q0PMG2) was inserted in-frame in a modified pCMV6-XL4 expression vector containing a leader peptide (PLP, prolactin leader peptide) followed by a N-terminal FLAG tag, MDGA1 insert, a 3CPro cleavage site and the human Fc domain. Secreted dimeric C-terminally Fc-tagged MDGA1 stably expressed in HEK293T cells was collected in serum-free Opti-MEM (Thermo Fisher Scientific, Inc.). Fc-tagged MDGA1 protein was run on an affinity column packed with Protein-G Plus Agarose fast flow resin (Pierce) using a gravity-flow system. Affinity column was washed with 250 mL wash buffer (50 mM HEPES pH 7.4, 300 mM NaCl) and eluted with 10 mL IgG elution buffer (Pierce) per the manufacturer’s instructions. For non-Fc-tagged MDGA1 protein used for immunization, following passage of conditioned medium through the column packed with Protein-G Agarose, the column was washed with 250 mL wash buffer (450 mM NaCl, 50 mM Tris, 1 mM EDTA, pH 8.0), the Fc tag was cleaved by O/N incubation with GST-tagged 3C PreScission Protease (GE Healthcare) in cleavage buffer (150 mM NaCl, 50 mM Tris, 1 mM EDTA, 1 mM DTT, pH 8.0), and the cleaved protein was collected in the eluate. The protease was subsequently separated from the eluted proteins using a Glutathione Sepharose (GE Healthcare) packed column. Fc-tagged and non-Fc-tagged proteins were concentrated using Amicon Ultra 10 kDa MWCO centrifugal filter units (Millipore), dialyzed against PBS, and protein concentration determined by Bradford assay (Bio-Rad). Immunization of rabbits and harvesting of polyclonal antiserum was performed by Synaptic Systems (MDGA1 polyclonal antiserum #421 002).

### Immunohistochemistry

Vibratome sections (80 µm) from the brains of either adult wildtype mice or *Mdga1* KO mice ^36^ (a gift of T. Yamamoto, Kagawa University, Japan) were permeabilized at RT for 40 min in PBS containing 0.5% Triton X-100 (Sigma-Aldrich). Sections were then blocked overnight at 4°C in PBS containing 10% normal horse serum (NHS), 0.5% Triton X-100, 0.5M glycine (Sigma-Aldrich, #G8898), 0.2% gelatin (Sigma-Aldrich, #G7041). Sections were then washed in PBS-0.5% Triton X-100 at RT and incubated at 4°C for 48 h with MDGA1 antiserum (dilution 1:500) in PBS containing 5% NHS, 0.5% Triton X-100 and 0.2% gelatin. Afterwards, sections were washed in PBS-0.5%Triton X-100 at RT before overnight incubation at 4°C with Alexa555-conjugated donkey-anti-rabbit antibody (Invitrogen, #A32794) in PBS containing 5% NHS, 0.5% Triton X-100 and 0.2% gelatin. Before mounting coverslips with Mowiol-4-88 (Sigma-Aldrich), sections were washed in PBS-0.5% Triton X-100. Images were acquired using a Leica SP8 confocal microscope (Leica Microsystems).

### Rat hippocampal cultures and electroporation

Gestant Sprague-Dawley rat females were purchased from Janvier Labs (Saint-Berthevin, France). Animals were handled and killed according to European ethical rules. Dissociated neuronal cultures were prepared from E18 rat embryos or P0 mice as previously described ^64^. Dissociated cells were electroporated with the Amaxa system (Lonza) using 300,000 cells per cuvette. Depending on the experiments, the following plasmid combinations were used: 1/ Homer1c-DsRed: shMDGA1 or shMDGA2: AP-MDGA1 rescue or AP-MDGA2-rescue: BirA^ER^ (1:3:1:1 µg DNA); 2/ Homer1c-DsRed and GFP-GPI (1:1 µg DNA); 3/ Homer1c-DsRed plus shCTRL (shMORB), shMDGA1, or shMDGA2 (1:3 µg DNA); 4/ Homer1c-DsRed: shCTRL, shMDGA1 or shMDGA2: AP-NLGN1: BirA^ER^ (1:3:1:1 µg DNA); 5/ Homer1c-DsRed: BirA^ER^: shCTRL or shMDGA2: AP-NLGN1: HA-MDGA2 rescue (1:1:3:1:1 µg DNA); 6/ Xph20-mRuby2: CRISPR/Cas9 CONTROL, CRISPR/Cas9 MDGA1, or CRISPR/Cas9 MDGA2: HA-MDGA2 rescue (1:3:1 µg DNA). Electroporated neurons were resuspended in Minimal Essential Medium (Thermo Fisher Scientific, #21090.022) supplemented with 10% Horse serum (Invitrogen) (MEM-HS), and plated on 18 mm glass coverslips coated with 1 mg/mL polylysine (Sigma-Aldrich, #P2636) overnight at 37°C. Three hours after plating, coverslips were flipped onto 60 mm dishes containing 15 DIV rat hippocampal glial cells cultured in Neurobasal plus medium (Gibco Thermo Fisher Scientific, #A3582901) supplemented with 2 mM glutamine and 1x B27^TM^ plus Neuronal supplement (Gibco Thermo Fisher Scientific, #A3582801). Cells were cultured during 8-14 days at 37°C and 5% CO_2_. Astrocyte feeder layers were prepared from the same embryos, plated between 20,000 and 40,000 cells per 60 mm dish previously coated with 0.1mg/mL polylysine and cultured for 14 days in MEM containing 4.5 g/L glucose, 2 mM L-glutamax (Sigma-Aldrich, #3550-038) and 10% horse serum. Ara C (Sigma-Aldrich, #C1768) was added after 3 DIV at a final concentration of 3.4 µM.

### Genomic cleavage of CRISPR constructs

To validate the genomic cleavage, we used a T7 endonuclease based method (GeneArt^TM^ Genomic detection kit, Thermo Fisher Scientific, #A24372). Briefly, dissociated hippocampal neurons where electroporated as described above with the CRISPR/Cas9 constructs and plated on glass coverslips. After 10 DIV, neurons were scraped in PBS and centrifuged at 1000xg for 5 min. Cells where then resuspended in 50 µL of lysis buffer containing 2 µL of protein degrader to extract genomic DNA. Then, PCRs were run to amplify a 555 bp genomic segment for MDGA1 and 546 bp for MDGA2. The following pairs of primers were used: MDGA1: F: 5’GGGAAGAGGTAGAGACCCAAGT 3’ R: 5’CCTCCATCAACACATAACGAAA 3’. MDGA2: F: 5’GCTGATAGGGAAGGACAGACAG 3’; R: 5’TAAATCCAAGACTGCAAGAGCC 3’. After checking the presence of the PCR fragments in an agarose gel, 1 µL of PCR reaction was denatured, reannealed, and digested with T7 endonuclease to reveal the presence of mismatches in the annealed fragments. Cleavage bands were detected in agarose gels.

### RT-qPCR

RNA was extracted from Banker neuronal cultures using the QIAzol Lysis Reagent (Qiagen) and the Direct-Zol RNA microprep (Zymo Research, cat#R2062) per manufacturer’s instructions. cDNA was synthetized using the Maxima First Strand cDNA Synthesis kit (Thermo Fischer Scientific, # K1641). At least three neuronal cultures were analyzed per condition and triplicate qPCR reactions were made for each sample. Transcript-specific primers were used at 600 nM and cDNA at 10 ng in a final volume of 10 µL. The LightCycler 480 SYBR Green I Master qPCR kit (Roche) was used according to manufacturer’s instructions. The Ct value for each gene was normalized against that of Sdha and U6. The relative level of expression was calculated using the comparative method (2ΔΔCt) ^65^. The following set of primers were used: MDGA1 Forward: 5’ GTTCTACTGCTCCCTCAACC 3’ Reverse: 5’ CGTTACCTTTATTACCGCTGAG 3’ MDGA2 Forward: 5’ AAGGTGACATCGCCATTGAC 3’ Reverse: 5’ CCACGGAATTCTTAGTTGGTAGG 3’ U6 Forward: 5′ GGAACGATACAGAGAAGATTAGC 3′ U6 Reverse: 5′ AAATATGGAACGCTTCACGA 3′ SDHA Forward: 5′ TGCGGAAGCACGGAAGGAGT 3′ SDHA Reverse: 5′ CTTCTGCTGGCCCTCGATGG 3′.

### Culture and transfection of COS-7 cells

COS-7 cells (from ATCC) were cultured in DMEM (Eurobio) supplemented with 1% glutamax (Sigma-Aldrich, #3550-038), 1% sodium pyruvate (Sigma-Aldrich, #11360-070), 10% Fetal Bovine Serum (Eurobio). For streptavidin pull-down and Western blots, COS-7 cells were plated in 6-well plates (100,000 cells/well) and transfected the next day using X-tremeGENE™ 9 DNA (Transfection Reagent, Roche), with HA-NLGN1 + AP-MDGA1 or AP-MDGA2 + BirA^ER^ (1 μg/well). Cells were left under a humidified 5% CO_2_ / 37°C atmosphere for 2 days before being processed for immunoprecipitation. For imaging experiments and shRNA validation experiments, cells were electroporated with the Amaxa system (Lonza) using the COS-7 ATCC program. Typically, 500,000 cells were electroporated with: 2 µg HA-MDGA1 or HA-MDGA2: 2, 4, 6 µg shMDGA1 or shMDGA2: 2 µg HA-MDGA1 or HA-MDGA2 rescue. After 24 h, cells were processed for imaging or biochemistry.

### Neuronal lysates and brain tissue subcellular fractionation

For biochemistry experiments, hippocampal neurons were plated at a density of 500,000 cells per well in a 6-well plate previously coated with 1 mg/mL polylysine for 24 hr at 37°C. Cells were cultured for 7, 14 and 21 DIV in Neurobasal medium supplemented with 2 mM glutamine and 1x NeuroCult SM1 Neuronal supplement. After 3 DIV, Ara C was added to the culture medium at a final concentration of 3.4 µM. Before lysis, plates were rinsed once in ice cold PBS and then scraped into 100 µL of RIPA buffer (50 mM Tris-HCl pH 7.5, 1 mM EDTA, 150 mM NaCl, 1% Triton-X100) containing protease inhibitor Cocktail Set III (Millipore #539134). Homogenates were kept for 30 min on ice and then centrifuged at 8000xg for 15 min at 4°C to remove cell debris. Protein concentration was estimated using the Direct Detect® Infrared Spectrophotometer (Merck-Millipore). 100 µg protein where loaded on a gel to detect endogenous MDGA proteins in Western blots. For all other proteins, 20 µg were loaded. Rat brain subcellular fractionation was performed as previously described ^66^.

### NLGN immunoprecipitation in neuronal cultures

Dissociated cells were electroplated with the Amaxa system (Lonza) using 1.5 x 10^6^ cells per cuvette and 8μg of shCONTROL or shMDGA2. Electroporated cells were plated at a density of 500.000 cells per well in a 6 well plate. At DIV 10 cells were treated with 3 mM pervanadate for 15 min at 37°C before lysis, to preserve phosphate groups on NLGNs. Whole-cell protein extracts were obtained by solubilizing cells in lysis buffer (50 mM HEPES, pH 7.2, 10 mM EDTA, 0.1% SDS, 1% NP-40, 0.5% DOC, 2 mM Na-Vanadate, 35 µM phenylarsine oxide, 48 mM Na-Pyrophosphate, 100 mM NaF, 30 mM phenyl-phosphate, 50 µM NH_4_-molybdate, 1 mM ZnCl_2_) containing protease Inhibitor Cocktail Set III (Millipore #539134). Lysates were clarified by centrifugation at 8000×g for 15 min. For immunoprecipitations, 500–1000 µg of total protein (estimated by Direct Detect® Infrared Spectrophotometer assay, Merck Millipore), were incubated overnight with 2 µg of antibody raised against an intracellular epitope in mouse NLGN1 (aminoacids 826 to 843) and which recognizes all NLGNs 1/2/3/4 (Synaptic Systems, #129 213). Antibody-bound NLGNs were incubated for 1 hour with 20 µL of protein G beads (Dyna-beads Protein G, Thermo Fisher Scientific) precipitated and washed 4 times with lysis buffer. At the end of the immunoprecipitation, 20 µL beads were resuspended in 20 µL of Laemli Sample Buffer buffer 2X (Biorad, #1610747), and supernatants were processed for SDS-PAGE and Western blotting.

### Streptavidin pull-down

Biotinylated AP-tagged MDGA1 or NLGN1 expressing COS-7 cells were rinsed once in ice cold PBS and then scraped in 100 µL RIPA buffer (50 mM Tris-HCl pH 7.5, 1 mM EDTA, 150 mM NaCl, 1% Triton-X100) containing protease inhibitor cocktail (Millipore). Homogenates were kept for 30 min on ice, then centrifuged at 8000xg for 15 min at 4°C to remove cell debris. The supernatant was recovered and the protein concentration was estimated using the Direct Detect® Infrared Spectrophotometer (Merck-Millipore). 400 µg protein were incubated with 40 µL of streptavidin coupled Dynabeads^TM^ M-280 (Thermo Fisher Scientific, #11205D). After 1 hr of incubation at room temperature on a rotating wheel, tubes were placed in the magnetic column and the beads were washed three times with lysis buffer. Proteins where eluted from the beads by directly adding 20 µL of Laemli Sample Buffer buffer 2X (Biorad, #1610747). Samples where then processed for SDS-PAGE and Western blotting.

### SDS-PAGE and Western-Blotting

Samples were loaded in acrylamide-bisacrylamide 4-20% gradient gels (PROTEAN TGX Precast Protein Gels, BioRad) and run at 100 V for 1 hr. Proteins were transferred to a nitrocellulose membrane for immunoblotting using the TurboBlot system (BioRad). After 1 hr blocking with 5% non-fat dry milk in Tris-buffered saline Tween-20 solution (TBST: 28 mM Tris, 137 mM NaCl, 0,05% Tween-20, pH 7.4), membranes were incubated during 1 hr RT or overnight at 4°C, with the primary antibody diluted in TBST solution containing 1% dry milk: custom-made rabbit anti-MDGA1 (Synaptic Systems #421002), rabbit anti-HA (Cell Signaling #3724 (C29F4), 1:1000), rat anti-HA (Roche #1186742300, 1:1000), mouse anti-actin (Sigma-Aldrich #A5316, 1:10,000), mouse anti-GFP (Sigma-Aldrich #11814460001, 1:1000), rabbit anti-ßIII tubulin (Abcam #ab18207, 1:25,000), mouse anti-tubulin (Sigma-Aldrich #T4026, 1:5000), mouse anti-PSD95 7E3-1B8 (Thermo Scientific #MA1-046, 1:2000), mouse anti-Synaptophysin (SVP-38) (Sigma-Aldrich # S5768, 1:2000), mouse anti-pTyr (Cell Signaling #9411, 1:1000), rabbit anti-panNLGN (Synaptic Systems #129 213, 1:1000). After 3 washes in TBST, membranes were incubated with horseradish peroxidase-conjugated donkey anti-mouse or anti-rabbit secondary antibodies (Jackson Immunoresearch, #715-035-150 and #711-035-152, respectively, concentration: 1:5000) or fluorophore-conjugated goat anti-mouse or anti-rabbit secondary antibodies (IRDye 680LT anti-rabbit #926-6821, IRDye 680LT anti-mouse #926-68020, IRdye-800CW anti-rabbit #926-32211, IRdye-800CW anti-mouse #926-32210 LI-COR) for 1 hr at room temperature. Target proteins were detected by chemiluminescence with Clarity^TM^ Western ECL Substrate (Bio-Rad #170-5061) on the ChemiDoc Touch System (BioRad) or Odyssey Fc Imaging System (LI-COR) for fluorescent secondary antibodies. For quantification of band intensities, images were processed with the Gels tool of ImageJ. Normalization of protein loading was done using endogenous actin or tubulin present in the samples.

### Immunocytochemistry

To visualize endogenous MDGA1 proteins and AMPA receptors at the cell surface, neurons were incubated live for 10 min at 37°C with the respective antibodies (rabbit anti-MDGA1, Synaptic Systems #421002 1:50; rabbit anti-GluA1, Agrobio, clone G02141, 0.2 mg/mL−1, 1:50; mouse anti-GluA2, clone 15F1, gift from E. Gouaux, 1:200), all diluted in Tyrode solution (15 mM D-glucose, 108 mM NaCl, 5 mM KCl, 2 mM MgCl_2_, 2 mM CaCl2 and 25 mM HEPES, pH = 7.4, 280 mOsm) containing 1% BSA. Then, cultures were fixed for 15 min in 4% paraformaldehyde, 4% sucrose, quenched in NH_4_Cl 50 mM in PBS for 15 min, permeabilized for 5 min with 0.1% Triton X-100 in PBS. After blocking during 20 min in PBS containing 1% BSA, cells were counter-stained for pre- and post-synaptic markers with a mixture of the following primary antibodies: anti-PSD-95 (Thermo Fisher Scientific #MA1-046, 1:100) and anti-VGLUT1 (Merck Millipore, #AB5905, 1:2000). Following 3 washes in PBS, cells were incubated with appropriate secondary antibodies coupled to Alexa fluorophores (405, 488, 564, or 647) (Thermo Fisher Scientific).

MDGA1 immunostainings were visualized on a commercial Leica DMI6000 TCS SP5 microscope using a X63, 1.4 NA oil objective and a pinhole opened to one Airy disk. Images of 1024×1024 pixels were acquired at a scanning frequency of 400 Hz. All other immunostainings were visualized using an inverted epifluorescence microscope (Nikon Eclipse TiE) equipped with a 60x/1.45 NA objective and filter sets for BFP (Excitation: FF02-379/34; Dichroic: FF-409Di03; Emission: FF01-440/40); EGFP (Excitation: FF01-472/30; Dichroic: FF-495Di02; Emission: FF01-525/30); Alexa568 (Excitation: FF01-543/22; Dichroic: FF-562Di02; Emission: FF01-593/40); and Alexa647 (Excitation: FF02-628/40; Dichroic: FF-660Di02; Emission: FF01-692/40) (SemROCK). Images were acquired with an sCMOS camera (PRIME 95B, Photometrics) driven by the Metamorph® software (Molecular Devices). The number of PSD-95 and VGLUT1 puncta per dendrite neuron length was measured using a custom macro written in Metamorph. Briefly, epifluorescence images of pre- and post-synaptic markers where first thresholded and segmented using the morphometric image analysis module of MetaMorph for structures bigger than 4 pix² (0.137 μm^2^). Then, the total length of the dendrite was measured with the free line drawing tool of MetaMorph, and the linear pre- and postsynaptic density was calculated.

### Single molecule tracking (uPAINT)

Universal Point Accumulation in Nanoscale Topography (uPAINT) experiments were performed as previously described ^21^. In brief, neuronal cultures were placed in a Inox Ludin chamber (Life Imaging Services) containing pre-warmed Tyrode solution supplemented with 1% biotin-free BSA (Roth #0163.4, Germany). The chamber was placed on a motorized inverted microscope (Nikon Ti-E Eclipse) enclosed in a thermostatic box (Life Imaging Services) providing air at 37°C. Biotinylated AP tags in MDGA1, MDGA2 and NLGN1 were labelled with STAR 635P-conjugated mSA at a concentration of 1 nM; N-terminal V5 tags in MDGA1, MDGA2 and LRRTM2 were labelled with 1 nM of recombinant Fab fragment coupled to STAR 635P (Abnova, #RAB00032). GFP-GPI was labeled with 1nM anti-GFP nanobody coupled to Atto647N. Endogenous AMPA receptors were labelled with a low concentration (∼1nM)of Atto 647N-conjugated anti-GluA2 antibodies. A four-color laser bench (405; 491; 561; and 647 nm, 100 mW each; Roper Scientific) is connected through an optical fiber to the TIRF illumination arm of the microscope and laser powers are controlled through acousto-optical tunable filters driven by Metamorph. The fluorophores STAR 635P and Atto 647N were excited with the 647-nm laser line through a four-band beam splitter (BS R405/488/561/635, SemRock). Samples were imaged by oblique laser illumination, allowing the excitation of individual fluorescent probes (mSA, V5 Fab, anti-GluA2) bound to the neuron surface, with minimal background coming from the probes in solution. Fluorescence light was collected through a 100 X/1.49 NA PL-APO objective using a FF01-676/29 nm emission filter (SemRock), placed on a filter wheel (Suter). Image stacks of 2,000–4,000 consecutive frames with an integration time of 20 ms, were acquired with a EMCCD camera working at 10 MHz and Gain 300 (Evolve, Photometrics, USA).

### dSTORM

AP-tagged proteins were labelled for dSTORM using a high concentration (100 nM) of mSA-Alexa647, in Tyrode solution containing 1% biotin free-BSA (Roth #0163.4, Germany) for 10 min at 37°C. V5-tagged proteins were labelled using 100 nM Alexa 647-conjugated anti-V5 Fab. GFP-GPI was labeled using 100nM anti-GFP nanobody coupled to Alexa647. Cells were rinsed and fixed with 4% PFA–0.2% glutaraldehyde in PBS-sucrose 4% for 10 min at room temperature, then kept in PBS at 4°C until dSTORM acquisitions. Neurons were imaged in Tris-HCl buffer (pH 7.5), containing 10% glycerol, 10% glucose, 0.5 mg/mL glucose oxidase (Sigma), 40 mg/mL catalase (Sigma C100-0,1% w/v) and 50 mM β-mercaptoethylamine (MEA) (Sigma M6500) ^67^. The same microscope described above for uPAINT was used. This microscope is further equipped with a perfect focus system preventing drift in the z-axis during long acquisition times. Pumping of Alexa647 dyes into their triplet state was performed for several seconds using ∼60 mW of the 647 nm laser at the objective front lens. Then, a lower power (∼30 mW) was applied to detect the stochastic emission of single-molecule fluorescence, which was collected using the same optics and detector as described above. Multicolour Tetraspec fluorescent 100-nm beads (Invitrogen, #T7279) or nano-diamonds (Adamas Nanotechnologies, Inc., #ND-NV140nm) were added to the sample for later registration of images and lateral drift correction. Single-molecule detection was performed online with automatic feedback control of the lasers using the WaveTracer module running in Metamorph, enabling optimal single-molecule density during the acquisition. Acquisition sequences of 64,000 frames were acquired in streaming mode at 50 frames per second (20-ms exposure time), thus representing a total time of 1280 s = 21 min.

### Offline single molecule detection, trajectory analysis, and image reconstruction

Analysis of the image stacks generated by uPAINT and dSTORM was made offline under Metamorph, using the PALM-Tracer program based on wavelet segmentation for single molecule localization and simulated annealing algorithms for tracking ^68, 69^. For the analysis of uPAINT experiments, the instantaneous diffusion coefficient, D, was calculated for each trajectory from linear fits of the first 4 points of the mean square displacement (MSD) function versus time, for trajectories containing at least 10 points. For very confined trajectories, the fit of the MSD function can give negative values for diffusion coefficients: in that case, D is arbitrarily set at 10^-5^ µm²/s. The uPAINT sequences were also represented as density maps integrating all individual molecule detections. These super-resolved images were constructed using a zoom factor of 5, i.e. with a pixel size of 32 nm which is five times smaller than that of the original image (0.16 µm) and corresponds to the pointing accuracy of our system. To sort individual trajectories among synaptic and extra-synaptic compartments, post-synapses were identified by wavelet-based image segmentation ^70^ of the Homer1c-DsRed signal, and the corresponding binary masks were transferred to the single-molecule images for analysis. Synaptic coverage was determined from super-resolved detection maps as the ratio between segmented areas containing detections over the whole synaptic region determined from the low resolution Homer1c-DsRed image. dSTORM stacks were analyzed using the PALM-Tracer program, allowing the reconstruction of a unique super-resolved image of 32 nm pixel size (zoom 5 compared to the original images) by summing the intensities of all localized single molecules (1 detection per frame is coded by an intensity value of 1). The localization precision of our imaging system in dSTORM conditions is around 60 nm (FWHM) ^71^. For the analysis protein enrichment at post-synapses, the average number of detections within Homer1c puncta was divided by the the average number of extra-synaptic detections, both normalized per unit area.

### Electrophysiology

Electrophysiological recordings were carried out at room temperature on primary hippocampal neurons expressing CRIPSR/Cas9 and either control, MDGA1, or MDGA2 gRNAs. Neurons cultured on 18 mm coverslips were observed with an upright microscope (Nikon Eclipse FN1) equipped with a motorized 2D stage and micromanipulators (Scientifica). Whole-cell patch-clamp was performed using micropipettes pulled from borosilicate glass capillaries (Clark Electromedical) using a micropipette puller (Narishige). Pipettes had a resistance in the range of 4–6 MΩ. The recording chamber was continuously perfused with aCSF containing (in mM): 130 NaCl, 2.5 KCl, 2.2 CaCl_2_, 1.5 MgCl_2_, 10 D-glucose, 10 HEPES, and 0.02 bicuculline (pH 7.35, osmolarity adjusted to 300 mOsm), while the internal solution contained (in mM): 135 Cs-MeSO_4_, 8 CsCl, 10 HEPES, 0.3 EGTA, 4 MgATP, 0.3 NaGTP, and 5 QX-314. Salts were purchased from Sigma-Aldrich and drugs from Tocris. Neurons were voltage-clamped at a membrane potential of −70 mV and AMPA receptor-mediated mEPSCs were recorded in the presence of 0.5 μM TTX. We verified that CNQX (20 μM) blocked the recorded currents.

### Statistics

Statistical values are given as mean ± s.e.m., unless otherwise stated. Statistical significance was calculated using GraphPad Prism 8.0 (San Diego, CA). For most experiments, data did not pass the D’Agostino and Pearson tests for normality, so comparisons were made using the non-parametric Mann–Whitney test. For data sets containing more than two conditions, comparisons were made by one-way analysis of variance (ANOVA) with the Kruskal-Wallis test for non-parametric samples, followed by a post hoc multiple comparison Dunn’s test. The number of experiments performed and the number of cells examined are indicated in each figure.

## Supplementary Figures

**Supplementary Fig. 1.**
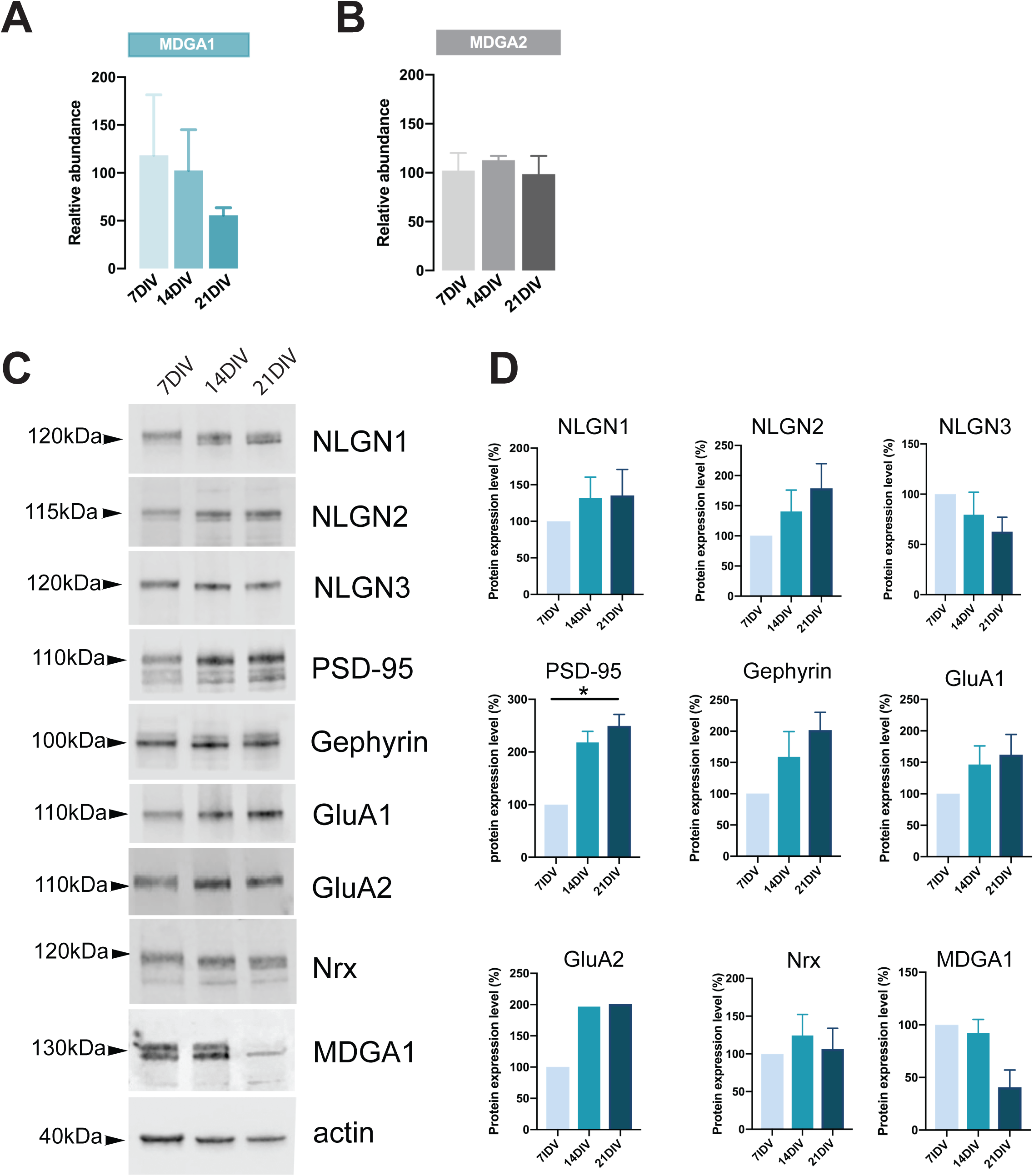
RT-qPCR evaluation of MDGA1 and MDGA2 mRNA expression levels and Western blot evaluation of protein expression during *in vitro* differentiation of hippocampal neurons. Dissociated hippocampal were cultured for 7, 14 and 21 DIV. **(A, B)** Normalized mRNA levels of MDGA1 (A) and MDGA2 (B) at different developmental stages, as determined by RT-qPCR. The Ct value for each gene was normalized against that of SDHA and U6 housekeeping genes, and expressed relatively to the value at 7 DIV. **(C)** Western-blots performed on protein extracts from hippocampal cultures, for different synaptic proteins. **(D)** Protein expression evaluation for the proteins detected in (C). For each protein the values are expressed in reference to the amount of protein detected at 7 DIV. All protein contents were normalized to actin.

**Supplementary Fig. 2.**
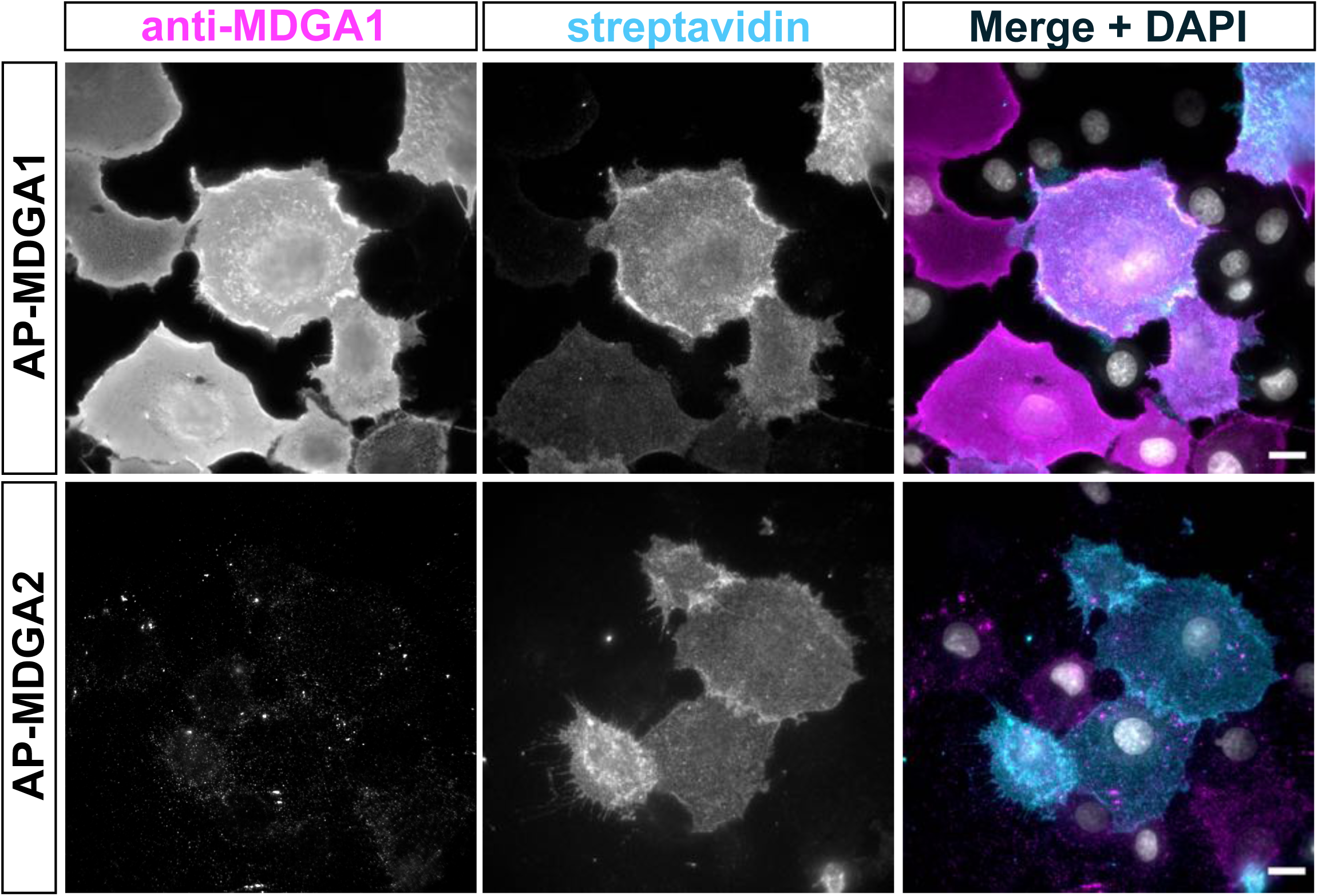
Surface labeling of COS-7 cells expressing recombinant MDGA1 or MDGA2 with the MDGA1 antiserum. COS-7 cells were co-electroporated with AP-MDGA1 or AP-MDGA2 and BirA^ER^. Cells were live labelled with anti-MDGA1 antibody and Alexa647-conjugated streptavidin. Following fixation, secondary anti-rabbit antibodies conjugated to Alexa546 were applied and images were acquired in the Alexa546 and Alexa647 channels. Merge images show anti-MDGA1 labelling in magenta, streptavidin in cyan, and DAPI staining in white. Note that the MDGA1 antibody recognizes only AP-MDGA1, and not AP-MDGA2. Scale bars, 10 µm.

**Supplementary Fig. 3.**
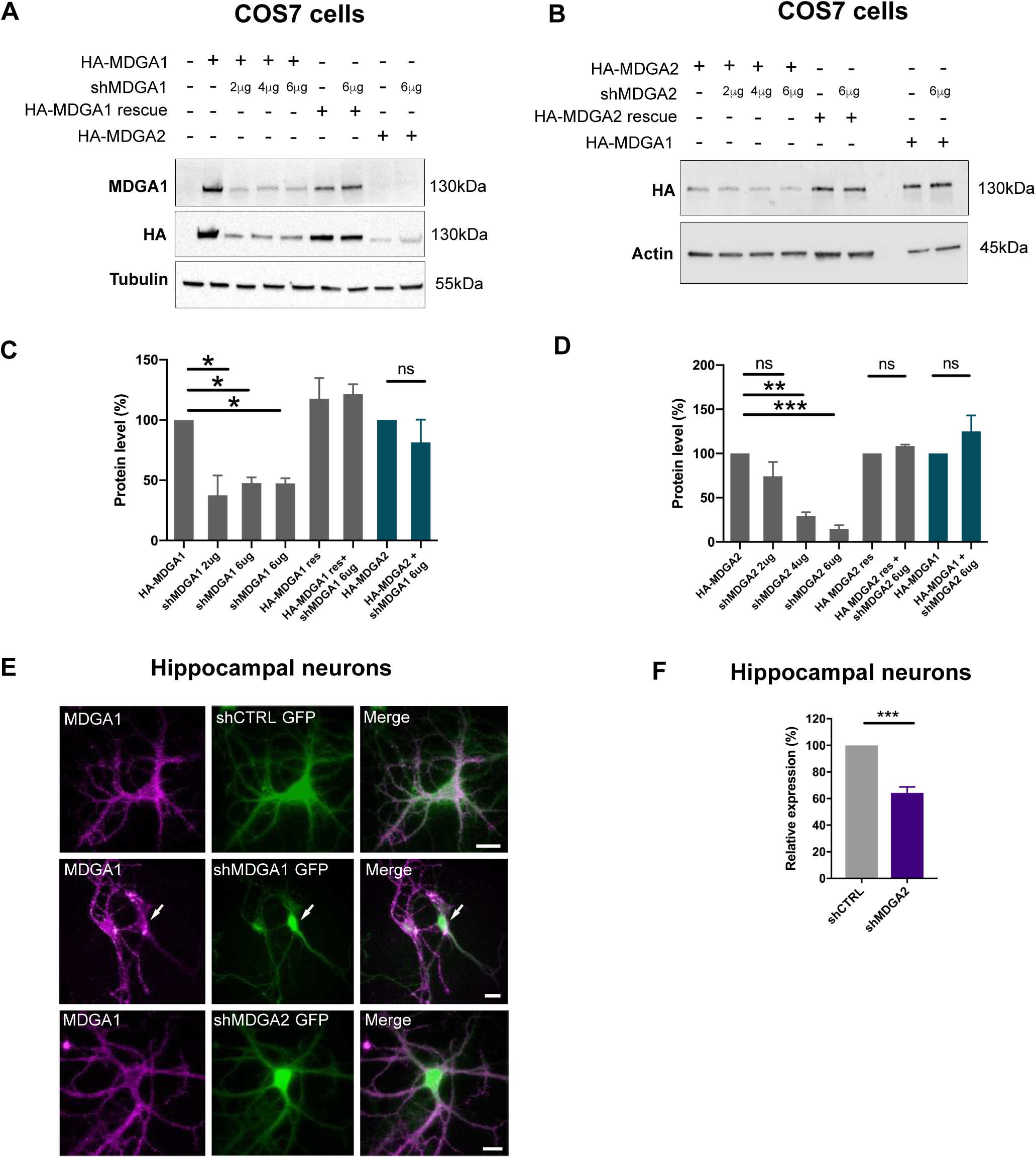
Validation of shRNA and rescue MDGA constructs in COS-7 cells and neurons. **(A)** Western-blots performed on protein extracts from COS-7 cells expressing HA-MDGA1, HA-MADGA1 rescue, or HA-MDGA2 with various doses of shMDGA1 (2, 4, and 6 µg). Blots were probed using antibodies to MDGA1, HA, and tubulin as a loading control. **(B)** Western-blots performed on protein extracts from COS-7 cells expressing HA-MDGA2, HA-MADGA2 rescue, or HA-MDGA1 with various doses of shMDGA2. Blots were probed using antibodies to HA or actin as a loading control. **(C, D)** Plots showing the quantitation of MDGA1 or MDGA2 levels normalized by tubulin or actin, and expressed in reference to the HA-MDGA1 control with no shMDGA1, or to HA-MDGA2 with no shMDGA2, respectively. Data represent mean ± SEM from 2 independent experiments, and were compared by Kruskal-Wallis test followed by Dunńs multiple comparison test (*P < 0.05; **P < 0.01; ***P < 0.001). **(E)** Immunodetection of endogenous MDGA1 at 14 DIV in neurons electroporated with shCTRL, shMDGA1 or shMDGA2 at DIV 0. Images show MDGA1 in magenta and the GFP reporter of all shRNAs in green. **(F)** RT-qPCR of MDGA2 mRNAs obtained from hippocampal neuronal cultures that were electroporated at DIV 0 with shCTRL or shMDGA2. PCR values were first normalized against U6 and SDHA housekeeping genes, and MDGA2 expression levels were then expressed as a function of shCTRL (taken as 100%). Data represent mean ± SEM from 4 independent experiments, and were compared by unpaired t-test (***P < 0.001).

**Supplementary Fig. 4.**
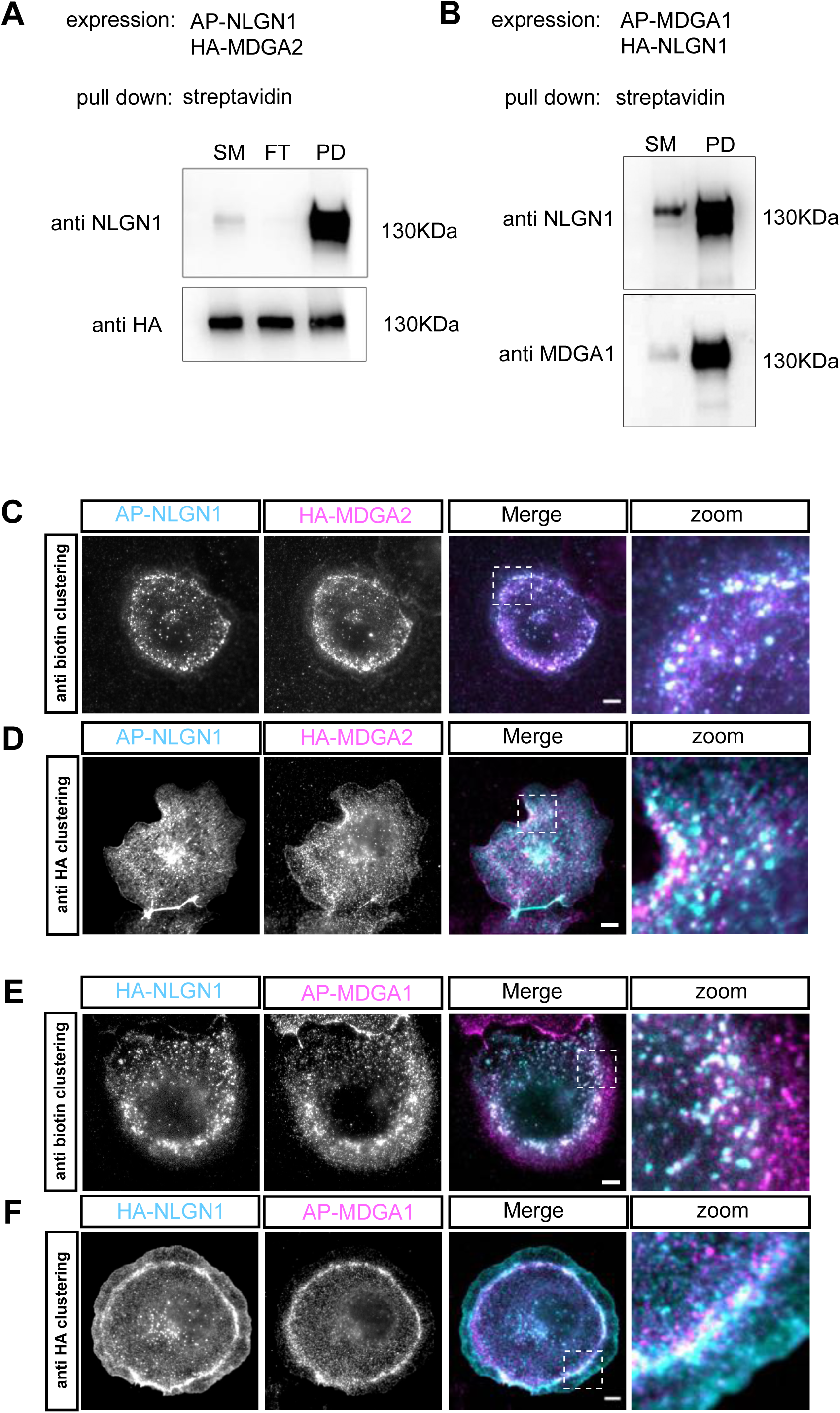
The labeling strategy does not impair the NLGN1-MDGA interaction. **(A)** COS-7 cells were co-transfected with AP-NLGN1, HA-MDGA2, and BirA^ER^, and biotinylated AP-NLGN1 was precipitated with streptavidin beads. Separated proteins were immunoblotted with anti-NLGN1 and anti-HA antibodies. SM: starting material, FT: flow through, PD: pull-down. **(B)** COS-7 cells were co-transfected with AP-MDGA1, HA-NLGN1, and BirA^ER^, and biotinylated AP-MDGA1 was precipitated with streptavidin beads. Separated proteins were immunoblotted with anti-NLGN1 and -MDGA1 antibodies. **(C, D)** Fluorescence microscopy of COS-7 cells expressing AP-NLGN1, BirA^ER^, and HA-MDGA2. **(C)** AP-NLGN1 was live clustered by incubating cells with a mix of primary anti-biotin antibody and an anti-mouse secondary antibody (cyan), and associated HA-MDGA2 was detected by rat anti-HA antibody followed by anti-rat antibody (magenta). Colocalization of NLGN1 and MDGA2 clusters indicates that accessibility of the AP tag is not impaired by the formation of NLGN1-MDGA2 complexes. **(D)** HA-MDGA2 was live clustered by cell incubation with a mix of rat anti-HA and secondary anti-rat antibody (magenta), then biotinylated AP-NLGN1 was detected with fluorescent streptavidin (cyan). Co-localization of NLGN1 and MDGA2 clusters was also observed. **(E, F)** COS-7 cells expressing AP-MDGA1, BirA^ER^, and HA-NLGN1. **(E)** AP-MDGA1 live clustered by incubating cells with a mix of primary anti-biotin antibody and anti-mouse secondary antibody (magenta), then HA-NLGN1 was detected by rat anti-HA antibodies followed by anti-rat antibody (cyan) **(F)** HA-NLGN1 was live clustered by incubating cells with a mix of anti-HA antibody and secondary antibody (cyan), then biotinylated AP-MDGA1 was detected with streptavidin (magenta). Co-localization of clusters was also observed. Scale bar, 10 µm.

**Supplementary Fig. 5.**
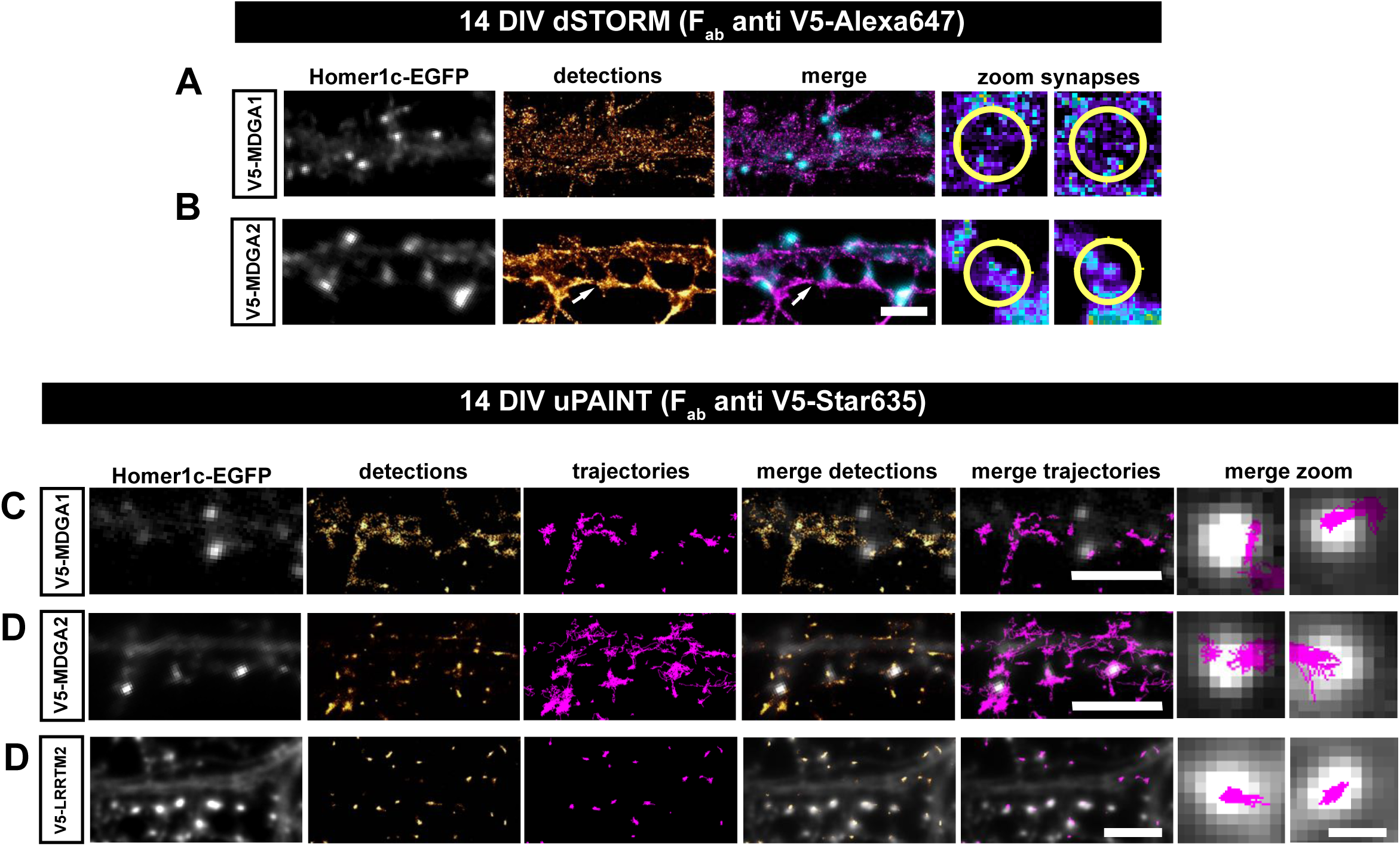
Lateral mobility and nanoscale localization of recombinant V5-MDGAs in hippocampal neurons. Dissociated rat hippocampal neurons were electroporated at DIV 0 with V5-MDGA1, V5-MDGA2, or V5-LRRTM2 as a positive control, together with the post-synaptic marker Homer1c-EGFP. **(A, B)** dSTORM images of MDGA1 and MDGA2 at the cell membrane, in DIV 14 neurons labeled with 100 nM Alexa647-conjugated anti-V5 Fab fragment. Representative images of dendritic segments showing Homer1c-EGFP positive synapses (grey), the super-resolved localization map of all V5-MDGA1 or V5-MDGA2 single molecule detections (gold), and merged images (Homer1c-EGFP in cyan and detections in magenta). Insets on the right show zoomed images of pseudo-colored localizations of V5-MDGAs in a synaptic area marked by a yellow circle. Arrows in B show an axon expressing V5-MDGA2 contacting spines in a dendrite also expressing V5-MDGA2. Scale bars, 5 µm. **(C, D, E)** uPAINT experiments were performed at DIV 14, after labelling neurons with STAR 635P-conjugated anti V5 Fab fragment Representative images of dendritic segments showing Homer1c-EGFP as synaptic marker (grey), the corresponding single molecule detections (gold) and trajectories (magenta). Superimposed images of synaptic markers and detections or trajectories are shown on the right of each panel with the same color code. Scale bars, 5 µm; 1 µm for insets.

**Supplementary Fig. 6.**
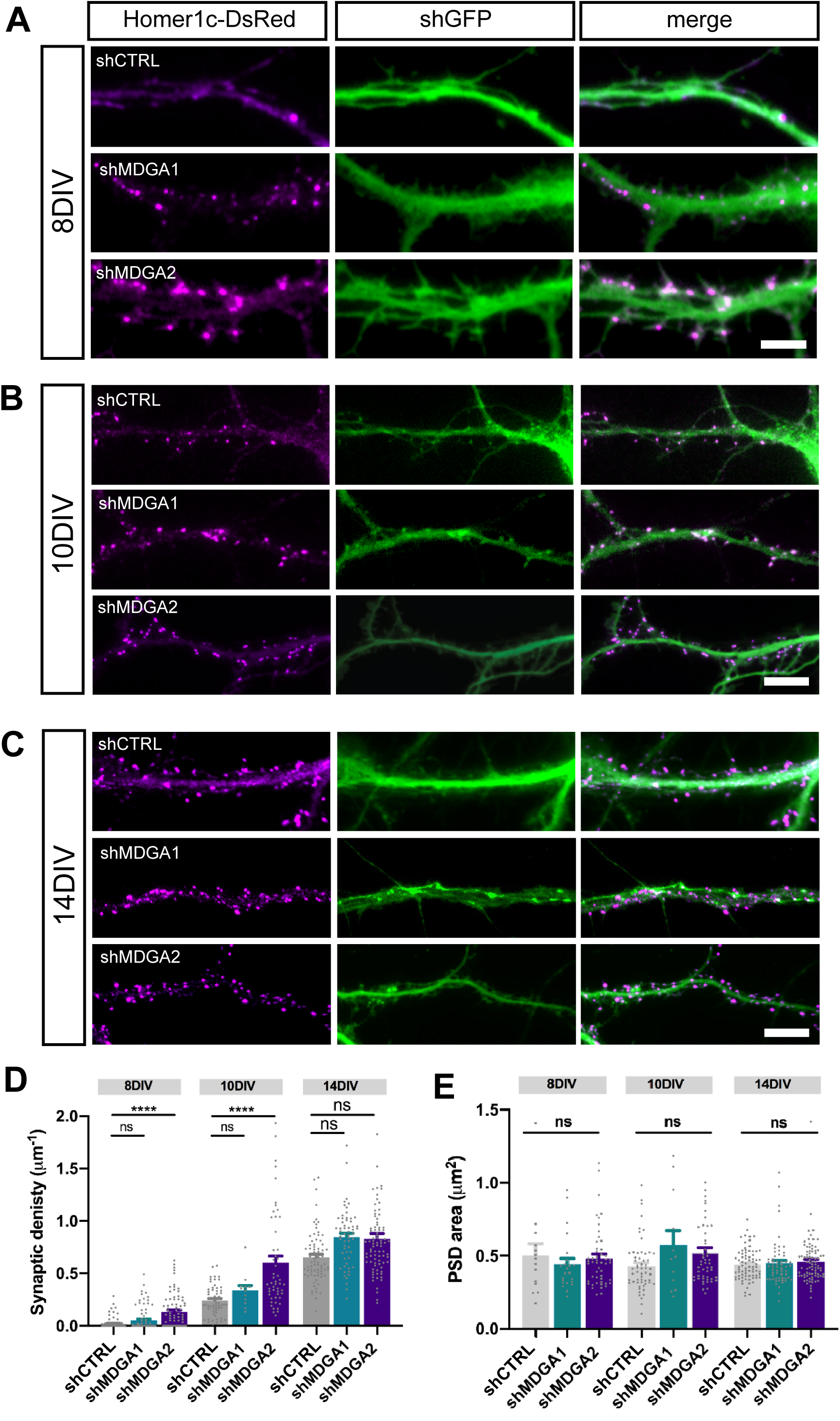
MDGAs knock-down increases synaptic density. Dissociated neurons where electroporated at DIV 0 with either shCTRL, shMDGA1, or shMDGA2. 8, 10 or 14 DIV after plating, epifluorescence images where acquired. **(A, B, C)** Representative images of dendritic segments at DIV 8, 10 and 14, respectively, showing Homer1c-DsRed (magenta), the GFP shRNA reporter (green), and merged images, at the different developmental stages. Scale bars, 5 μm (A), 10 μm (B, C). **(D, E)** Bar plots showing the density and area of individual Homer1c-DsRed puncta in the different developmental stages. Data represent mean ± SEM from at least three independent experiments, and were compared by a Kruskal–Wallis test followed by Dunn’s multiple comparison test (****P < 0.0001).

**Supplementary Fig. 7.**
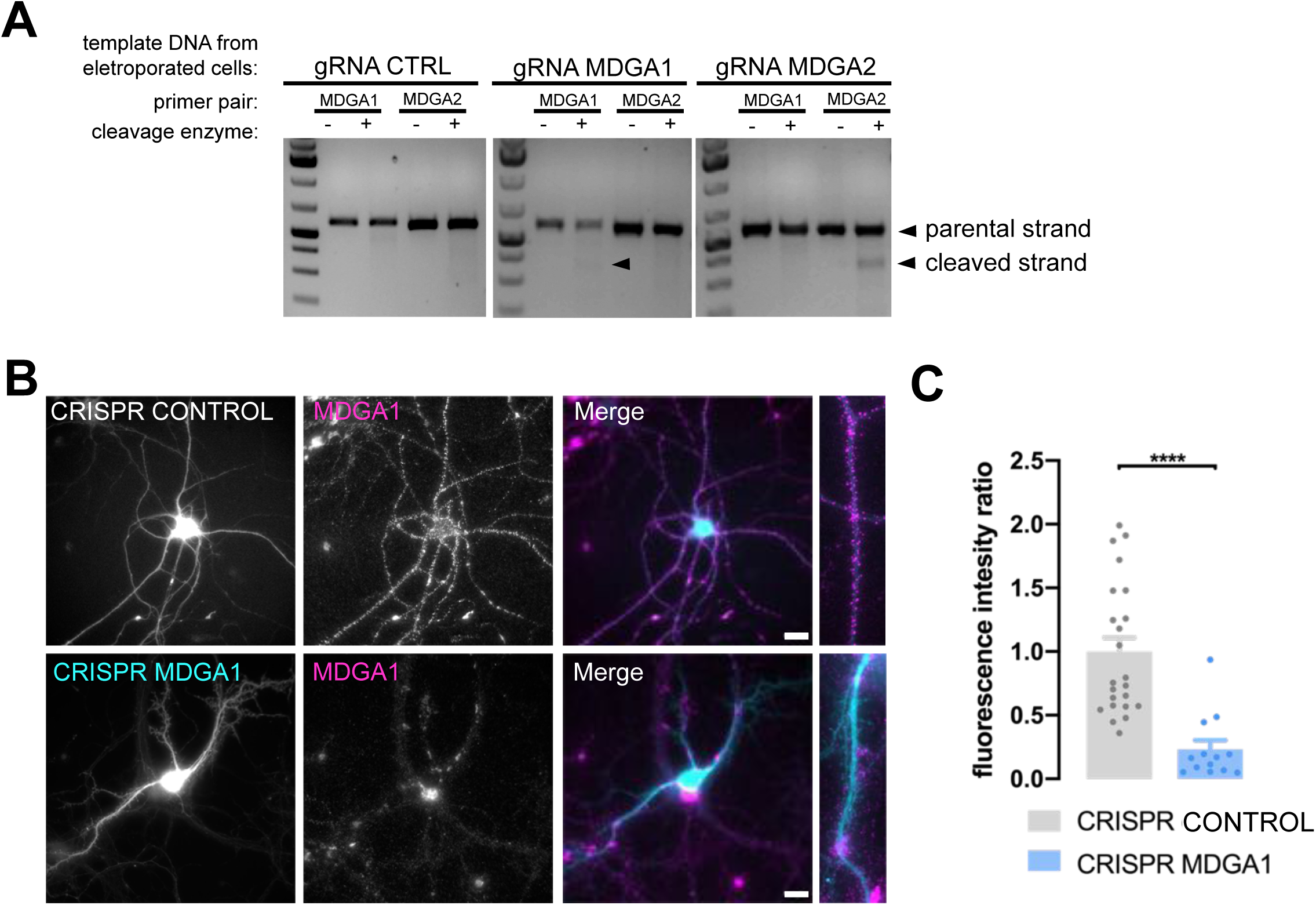
Validation of CRISPR/Cas9 knock-out of MDGA1/2. Hippocampal neurons were electroporated at DIV 0 with CRISPR/Cas9 control, CRISPR/Cas9 MDGA1, or CRISPR/Cas9 MDGA2**. (A)** At 10 DIV, genomic DNA was extracted and a T7 endonuclease based method was used to detect genomic cleavage in the CRISPR/Cas9 system. Cleavage bands were observed only for gRNA MDGA1 and gRNA MDGA2 when primers to amplify the target sequence were used and in the presence of T7 endonuclease. **(B)** Neurons were live immunolabelled with anti MDGA1 antibody at DIV 10. Scale bars, 10 µm. **(C)** Bar graph of fluorescence intensity ratio of MDGA1. Fluorescent results are expressed relative to CRISPR CONTROL values. Data represent mean ± SEM from two independent experiments (n > 10 neurons for each experimental condition). Values were compared by Mann-Whitney test (****p < 0.0001).

**Supplementary Fig. 8.**
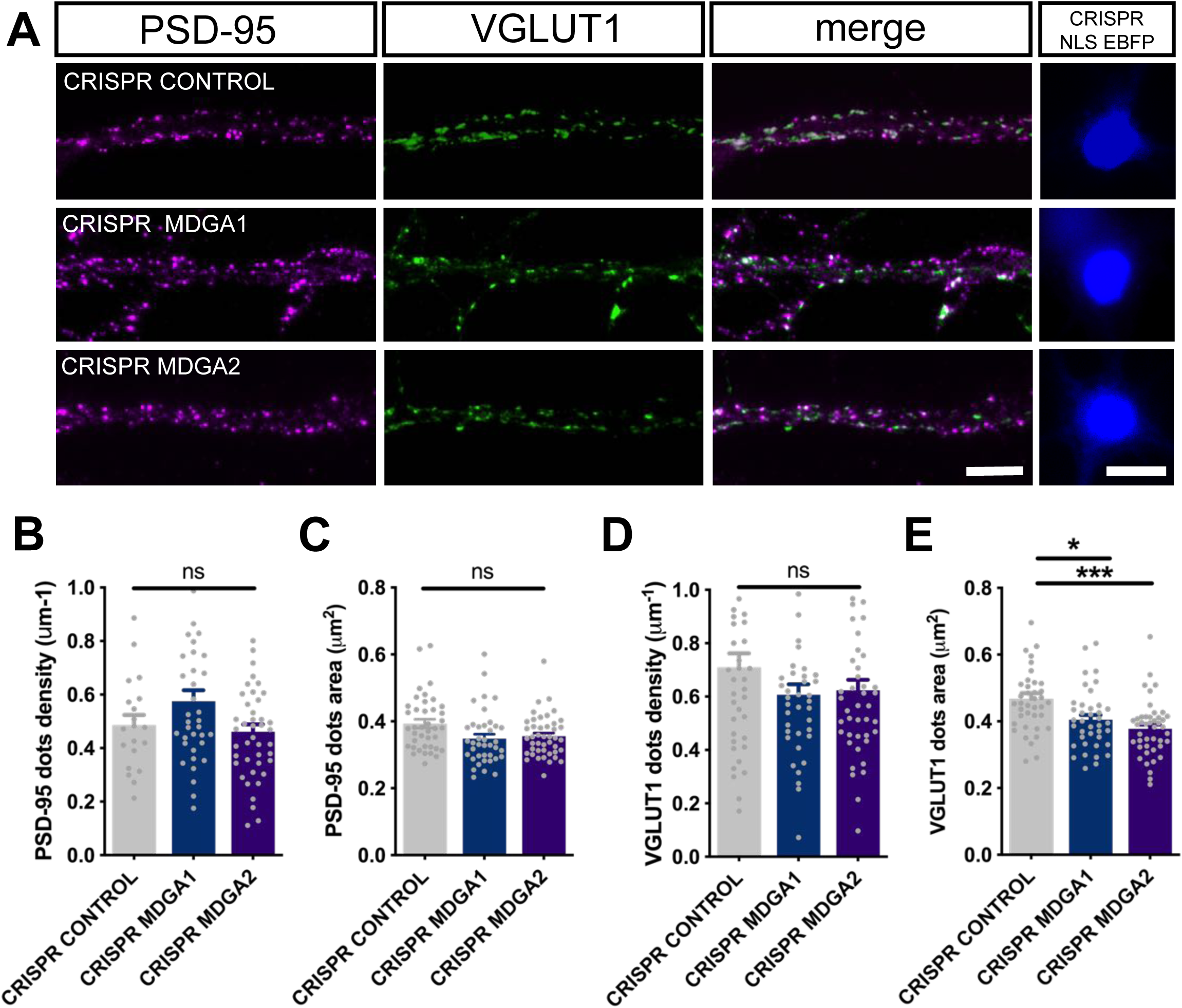
MDGAs knock-out has no effect in DIV 14 neurons. Dissociated neurons where electroporated at DIV 0 with either CRISPR/Cas9 control, CRISPR/Cas9 for MDGA1, or CRISPR/cas9 for MDGA2. 14 DIV after plating, cultures were fixed, permeabilized, and endogenous PSD-95 and VGLUT1 were immunostained. **(A)** Representative images of dendritic segments showing PSD-95 staining (magenta), VGLUT1 staining (green), the merged images, and the nuclear EBFP control of CRISPR/cas9 construct expression. Scale bars, 10 µm. **(B-E)** Bar plots showing the density and area of individual PSD-95 (B, C) and VGLUT1 (D, E) puncta in the different conditions. Data represent mean ± SEM from at least three independent experiments, and were compared by a Kruskal–Wallis test followed by Dunn’s multiple comparison test (*P < 0.05; ***P < 0.001).

**Supplementary Fig. 9.**
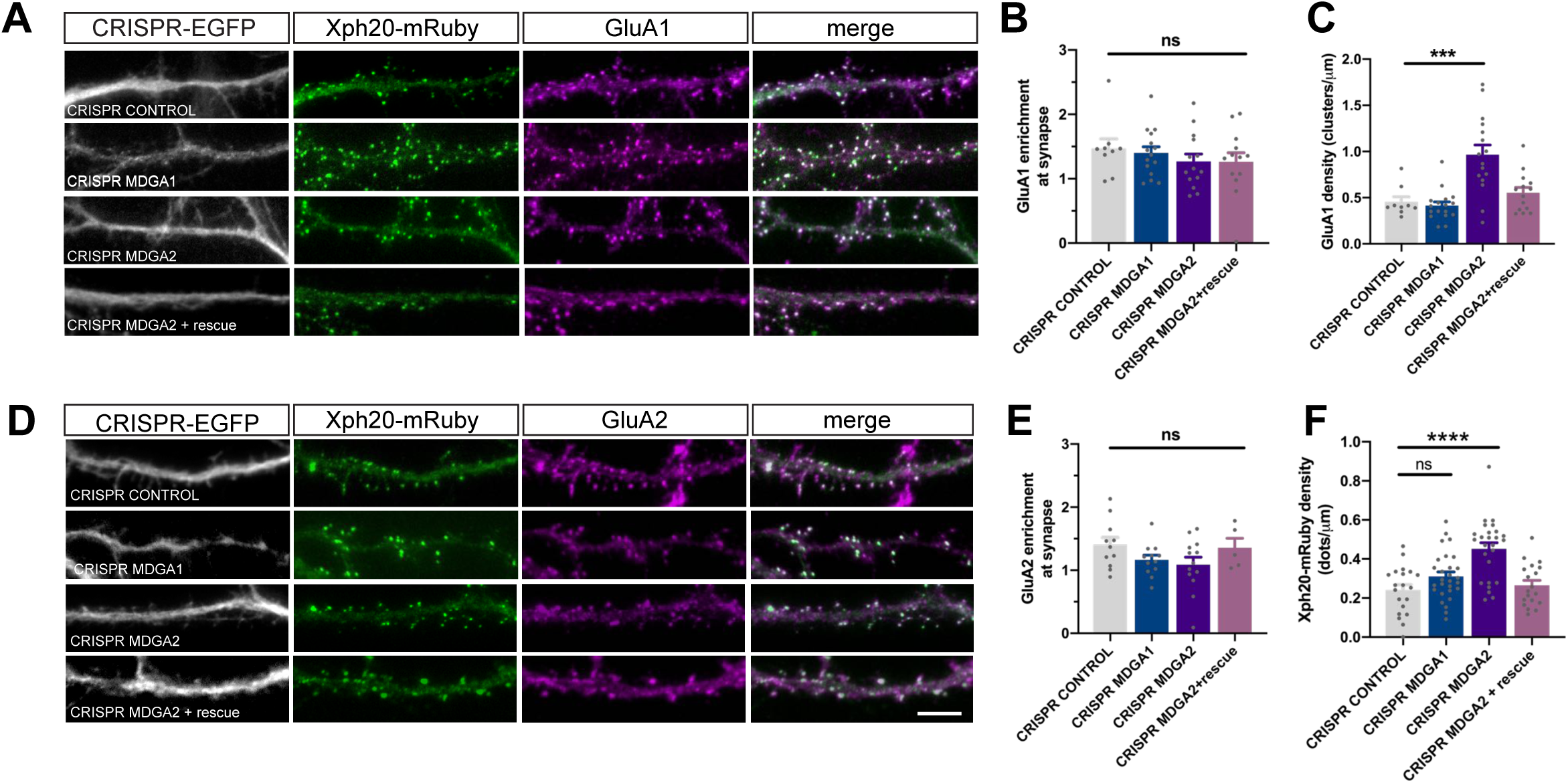
AMPA receptor membrane localization upon MDGA1/2 knockout. **(A, D)** Live labelling of GluA1 and 2 with specific antibodies (magenta) in CRISPR/Cas9 (white) and xph20-mRuby2 (green) expressing cells. Scale bar 10μm. **(B, E)** Bar plots representing the enrichment of GluA1 or GluA2 at the synapse, respectively, was evaluated measuring the fluorescence intensity of GluA1 or GluA2 at Xph20 positive sites and normalized by the fluorescence intensity of AMPA receptors at the dendritic shaft. **(C)** GluA1 cluster density in the dentrite was evaluated by thresholding the GluA1 signal and counting the segmented dots per unity of length. The same thresholding parameters were applied to all experimental conditions. **(F)** Xph20-mRuby2 density per dendrite unit length. Results represent mean ± SEM of at least 10 neurons per condition from two independent experiments, and were compared by Kruskal-Wallis test, followed by Dunn’s multiple comparison test (***P < 0.001; ****P < 0.0001)

